# Unveiling Mechanistic and Structural Insights of EstS1 Esterase: A Potent Broad-Spectrum Phthalate Diester Degrading Enzyme

**DOI:** 10.1101/2024.08.14.607922

**Authors:** Shalja Verma, Shweta Choudhary, K Amith Kumar, Jai Krishna Mahto, Anil Kumar Vamsi K, Ishani Mishra, Vellanki Bhanu Prakash, Debabrata Sircar, Shailly Tomar, Ashwani Kumar Sharma, Jitin Singla, Pravindra Kumar

## Abstract

The ubiquitous presence of plastics and plasticizers around the globe has raised an alarming condition. Phthalate diesters are high-priority pollutants that mimic natural hormones and act as endocrine disruptors upon entering living systems. While certain bacterial esterases have been identified for their role in phthalate diester degradation, their structural and mechanistic characteristics remain largely unexplored. A thermostable and pH-tolerant EstS1 esterase from *Sulfobacillus acidophilus* catalyzes the conversion of low molecular weight phthalate diesters to monoesters. This study highlights the unique potential of EstS1 to degrade high molecular weight bis(2-ethylhexyl) phthalate (DEHP) by employing biophysical and biochemical approaches along with in-depth structural analysis utilizing high-resolution crystal structures in both apo and complex forms, with various substrates, products, and their analogs to elucidate mechanistic details. The catalytic tunnel mediating entry and exit of the substrate and product, respectively, centralized the Ser-His-Asp triad performing catalysis by bi-bi ping-pong mechanism, forming a tetrahedral intermediate. Additionally, structural analysis of the polypropylene analog jeffamine with EstS1 revealed effective covalent binding, demonstrating its multifunctional capability. Mutation analysis showed that the Met207Ala mutation abolished DEHP binding at the active site, confirming its essential role in supporting catalysis. These findings underscore the potential of EstS1 as a key tool for advancing technologies aimed at phthalate diesters biodegradation.

## Introduction

Parallel to plastics, the environmental load of endocrine-disrupting, carcinogenic plasticizers, is increasing exponentially. Phthalate diester plasticizers non-covalently bind to plastics to impart durability and flexibility and often leach out of plastics in the environment (1). Some of them are released during the production and distribution process. Investigations have identified toxic levels of these plasticizers in body fluids and organs of different aquatic and terrestrial organisms including humans (2–4). Microbial biodegradation or bacterial assimilation of phthalate diesters utilizing complex enzymatic pathways is being investigated (Figure 1A) (5–13). Specific esterases or hydrolases for phthalate diester degradation have been identified in diverse microbial strains such as *Micrococcus sp.* YGJ1 (9), *Acinetobacter sp.* M673 (10), *Fusarium sps*., *Fusarium oxysporum f.sp*. pisi (11), *Gordonia sp.* strain 5F (7), Gordonia sp. strain P8219 (12) and *Sulfobacillus acidophilus* DSM10332 (13) etc. These esterases break down diesters into corresponding alcohols and phthalate monoesters (10, 12, 13) followed by dioxygenases, dehydrogenases, and decarboxylases ultimately leading to protocatechuate formation that eventually enters the tricarboxylic acid (TCA) cycle for complete metabolism (13–17).

**Figure 1:**
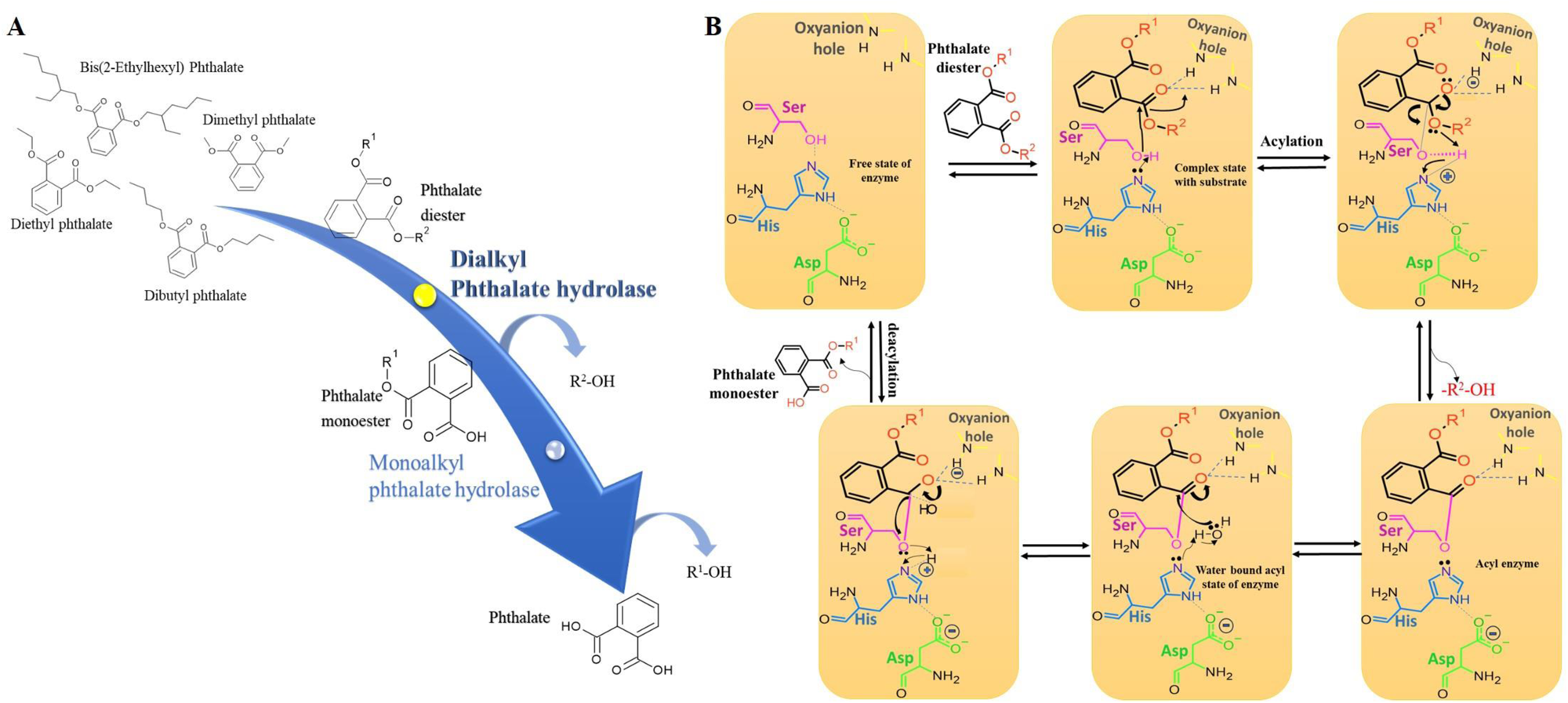
Enzymatic degradation of phthalate diesters. A) Dialkyl phthalate hydrolase degrade phthalate diesters to phthalate monoesters which is followed by degradation of phthalate monoesters to phthalate by monoalkyl phthalate hydrolase. B) Catalytic mechanism of phthalate diester degrading esterase or α/β hydrolase showing catalytic triad of Ser-His-Asp and arrangement of two amides (N-H’) of main chain forming oxyanion hole. The catalytic serine attack on the carbonyl carbon of the substrate and forms the 1^st^ tetrahedral intermediate. Histidine by acting as base deprotonates the catalytic serine. The alkyl alcohol group is released upon reformation of the double bond between the carbon and oxygen leading to acyl-enzyme intermediate. A water molecule enters and binds to the intermediate, now histidine by acting as base deprotonate this water molecule which attacks on acyl enzyme carbonyl carbon thus forming the 2^nd^ tetrahedral intermediate. Again, reformation of the double bond between carbon and oxygen releases the phthalic acid monoester releasing the free form of the enzyme.

Bis-(2-ethylhexyl) phthalate (DEHP), a lead priority pollutant, is a high molecular weight phthalate diester. In esters, the carbonyl dipole of the two ester dipoles (carbonyl and ether) contributes as a hydrogen acceptor via carbonyl oxygen thus, can convey water solubility. But this solubility is noticeably less than corresponding acids or alcohols as they are, incapable of both accepting and donating the hydrogen bonds to molecules of water, non-ionic in nature, and have bulky hydrocarbons (18). Similarly, in DEHP, this lack of solubility in water reduces its degradability and specific esterase enzymes that can effectively accommodate the bulky hydrocarbon groups of DEHP are needed. Enzymatic degradation of DEHP by esterase involves the carbonyl (C=O) carbon, due to its electrophilic nature, that is attacked by the nucleophilic catalytic residue of esterase forming a tetrahedral complex leading to cleavage of ester bond producing mono-(2-ethylhexyl) phthalate (MEHP) and 2-ethylhexanol (12, 13, 14, 18–25) (Figure 1B). Acidic and basic conditions enhance the carbonyl carbon electrophilicity and nucleophile concentration and thus increase the rate of catalysis. Esterases with wide pH tolerance can more efficiently catalyze the degradation of DEHP over a wide range of pH (26).

Several phthalate esters degrading esterases have been identified but only a few have been reported for degradation of DEHP. While these DEHP-degrading esterases have been biochemically analysed to provide insights into their degradation efficiency, thermostability, pH, and salt tolerance, the structural and mechanistic details remain elusive (7, 12, 14, 27, 28, 29). Understanding these details is crucial for enzyme engineering studies aimed at enhancing catalytic efficiency and expanding the range of substrates they can act on.

To fulfill this lacunae, a comprehensive biophysical, biochemical and in-depth structural characterization of thermostable and pH tolerant EstS1 esterase from *Sulfobacillus acidophilus* DSM10332 was conducted. A previous study reported its ability to degrade low molecular weight phthalate diesters (13). Here, we demonstrate that EstS1 also has significant potential to degrade the high molecular weight phthalate diester DEHP, thereby expanding its substrate range. High-resolution crystal structures of apo-EstS1 (1.22 and 1.55 Å) and complexes with various compounds, including DEHP reaction products (1.9 Å), dimethyl phthalate (1.6 and 1.5 Å), para-nitrophenyl butyrate (1.8 Å), monomethyl phthalate (2 Å), para-nitrophenol (1.5 Å), phthalate (1.5 Å), and jeffamine (1.5 Å), as well as the EstS1 active site mutant Ser154Ala with DEHP (2.4 Å), revealed the precise catalytic mechanism and the pathway for substrate entry and product exit through the enzyme’s catalytic tunnel. The strong covalent interaction with jeffamine, a polypropylene analog, observed in the crystal structure, indicated a high affinity of EstS1 for polypropylene derivatives at the active site. Site-directed mutagenesis studies showed that replacing the active site Ser154 with threonine slightly decreased DEHP binding, while replacing Met207 with alanine drastically reduced DEHP binding, highlighting the critical role of Met207 in substrate binding. This work opens up opportunities for further enzyme engineering studies aimed at enhancing the potential of EstS1 for developing engineered enzyme-based technologies for *in-situ* applications^37^.

## Results

### Purification of wild-type EstS1 and evaluation of its esterase activity by UV-spectroscopy

EstS1 (∼34KDa) wild-type was expressed in *E. coli* at optimized conditions for high protein yield using an IPTG (isopropyl β-D-1-thiogalactopyranoside) inducible pET28a vector. A highly purified (∼99% purity) monomeric form of EstS1 wildtype with N-terminal His_6_ tag using Ni-NTA chromatography was obtained at a concentration of 50mg/ml (Figure S1).

The catalytic activity of the EstS1 against para-nitrophenol butyrate (pNPB) revealed a specific activity of 338 U/mg of protein using 0.5 mM pNPB at 25 °C and pH 7.4. The enzyme kinetic parameters displayed K_m_ and V_max_ values of 0.18 mM and 451.95 μM/min/mg of protein respectively, which were consistent with the enzyme activity reported in a previous study conducted at 25 °C (13). Additionally, the catalytic efficiency, as indicated by the k_cat_/K_m_ value, was calculated to be 1398.72 μM^−1^ sec^−1^ (Figure S5).

### Fluorescence spectroscopy for DEHP binding analysis

Binding analysis of DEHP with EstS1 was conducted using fluorescence spectroscopy, with DEHP concentrations ranging from 1 μM to 5 μM while maintaining a constant concentration of EstS1 (2 μM). A proportional quenching of fluorescence was observed as the concentration of DEHP increased. Since the active site of EstS1 contains several tryptophan, phenylalanine, and tyrosine residues, the binding of DEHP at the active site significantly quenched the fluorescence intensity contributed by these amino acids (Figure 2A) (30).

**Figure 2:**
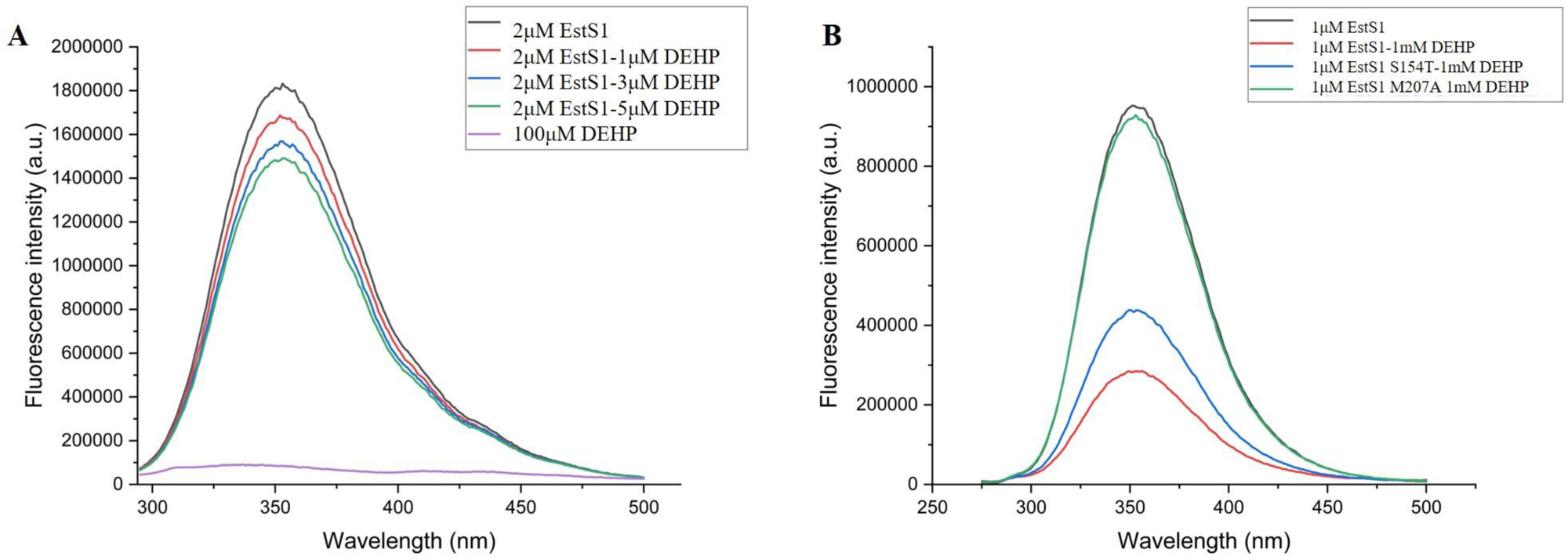
Fluorescence spectroscopy analysis of DEHP degradation by wild-type and mutant EstS1 esterase. A) Fluorescence intensity versus wavelength plots showing reduction in fluorescence intensity of EstS1 (2μM) when incubated with the increasing concentration of DEHP from 1μM to 5Μm. B) Fluorescence intensity versus wavelength plots showing variation in fluorescence intensity of wildtype and S154T and M207A mutant forms of EstS1 esterase when incubated with 1mM DEHP. The least reduction in intensity almost equivalent to EstS1 in absence of DEHP was observed in M207A EstS1 mutant with 1mM DEHP revealing effective role of M207 in mediating the binding of DEHP at the EstS1 catalytic site. The fluorescence intensity peak of S154T mutant showed less reduction in intensity compared to the wildtype form in presence of DEHP.

### Liquid Chromatography Mass Spectrometric (LC-MS) analysis of bis-(2-ethyl hexyl) phthalate (DEHP) degradation

The degradation of DEHP by EstS1 esterase into monoethylhexyl phthalate (MEHP) and 2-ethylhexanol was analyzed using LC-MS. A reaction mixture containing 10 μM EstS1 and 2 mM DEHP was continuously shaken at 200 rpm for 3 hours to increase entropy and, consequently, the reaction rate. Control samples of DEHP and MEHP without the enzyme were subjected to the same conditions for comparison. In the presence of EstS1, a decrease in DEHP concentration (retention time t_R_ = 1.73 min) was observed, indicated by a reduction in peak intensity compared to the control. Additionally, two new peaks appeared for MEHP at t_R_ = 2.93 min and 2-ethylhexanol at t_R_ = 1.40 min (Figure 3). The intensity versus m/z spectra at both low and high energies showed significant peaks at m/z ratios of 393, 277, and 132.96, corresponding closely to the exact molecular masses of 390.60, 278.34, and 130.23 g/mol for DEHP, MEHP, and 2-ethylhexanol, respectively (31).

**Figure 3:**
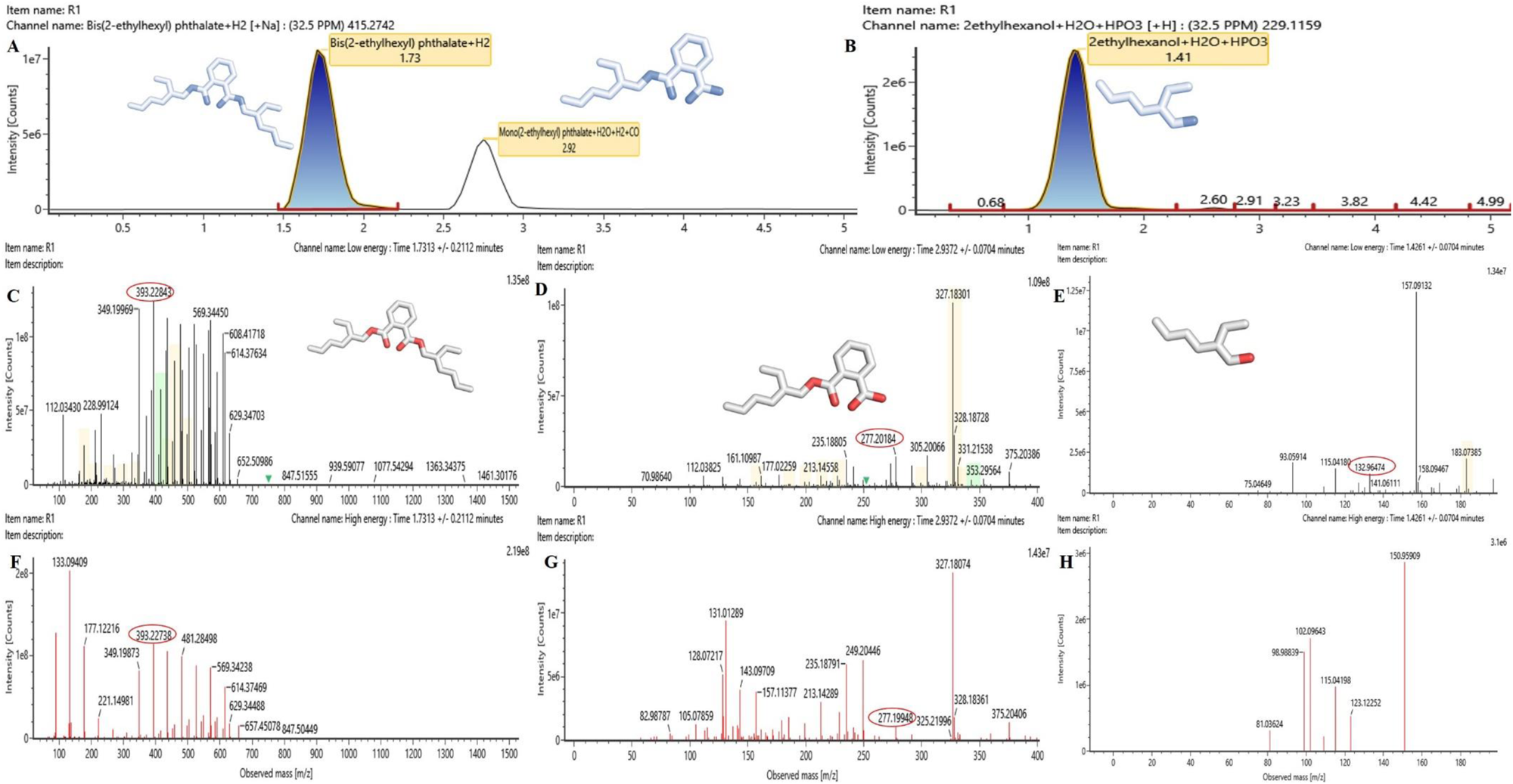
Liquid Chromatography Mass Spectrophotometry analysis of DEHP degradation by EstS1 esterase. A) Intensity versus retention time plot displaying intensity peaks of DEHP and MEHP at retention time of 1.73 and 2.92 min when EstS1 (10 μM) esterase was incubated with DEHP (2Mm) for 3hrs at 37 °C. B) Intensity versus retention time plot of 2-ethylhexanol a by-product of DEHP degradation at retention time of 1.41min. C) Intensity versus m/z ratio mass spectrum at low energy of DEHP showing significant peak at m/z ration of 393.22843 at retention time 1.73 min which is close to DEHP molecular weight of 390.55 Da. D) Intensity versus m/z ratio mass spectrum at low energy of DEHP degradation product MEHP showing significant peak at m/z ration of 277.20 at retention time 2.93 min which is close to MEHP molecular weight of 278.34 Da. E) Intensity versus m/z ratio mass spectrum at low energy of DEHP degradation product 2-ethylhexanol showing significant peak at m/z ratio 132.96 at retention time 1.42 min which is close to 2-ethylhexanol molecular weight of 130.229 Da. F) Intensity versus m/z ratio mass spectrum at high energy of DEHP showing significant peak at m/z ration of 393.227 at retention time 1.73 min G) Intensity versus m/z ratio mass spectrum at high energy of DEHP degradation product MEHP showing significant peak at m/z ration of 277.20 at retention time 2.93 min. H) Intensity versus m/z ratio mass spectrum at high energy of DEHP degradation product 2-ethylhexanol.

### High-performance liquid chromatography (HPLC) based DEHP degradation kinetics

To investigate the steady-state kinetics of DEHP degradation by EstS1 esterase, varying concentrations of DEHP were incubated with a constant concentration of EstS1 for 3 hours at 200 rpm. A standard plot of DEHP was established within the same concentration range for quantification. A prominent peak for DEHP at retention time t_R_ = 4.34 min was observed. The degradation of DEHP was quantified by the reduction in the peak area of DEHP peak in the presence of EstS1. The rate of degradation increased with increasing DEHP concentration until it reached a maximum velocity (Vmax) (Figure 4), after which saturation occurred, consistent with Michaelis-Menten kinetics (32, 33, 34). The V_max_ and K_m_ values for the DEHP degradation were determined and the catalytic efficiency (k_cat_/K_m_) was estimated to be 6.75 μM^−1^ sec^−1^. The degradation kinetics of DEHP by EstS1 indicate that the enzyme exhibits high efficiency compared to previous reports on degradation of phthalate diesters, including DEHP (7).

**Figure 4:**
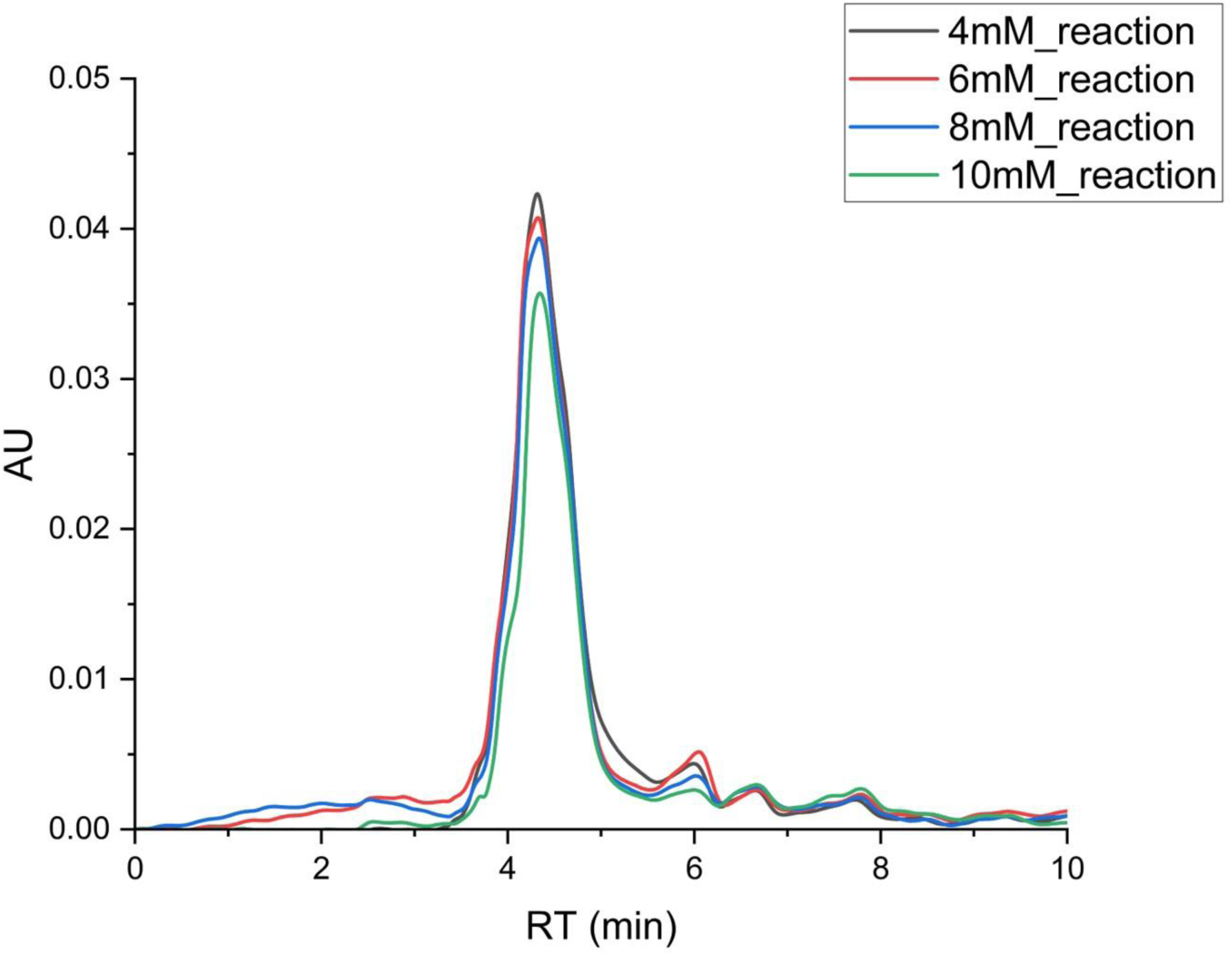
High Performance Liquid Chromatography analysis of DEHP degradation for kinetic analysis. Absorbance unit versus retention time plot at 254nm of different concentrations of DEHP incubated with EstS1 esterase revealed that the rate of reduction of absorbance of DEHP (indicating DEHP degradation) increases with increase in DEHP concentration in reaction, thus follows the first order Michaelis–Menten kinetics.

### X-Ray crystallography based structural study of EstS1 in apo and complex forms Structure of EstS1 esterase

The crystal structures of the apo-form of EstS1 were determined in the P63 space group with a single monomer in an asymmetric unit and parameters for refinement quality R*_work_* and R*_free_* were 17.30% and 20.40% for the structure at 1.22 Å resolution and 15.1% and 20.5% for the structure at 1.5 Å resolution, respectively (Table 1). The highly flexible loop region ranging from Pro16 to Gly22 was missing in the electron density map. The structure of EstS1 consists of 304 residues, and among them, 17 residues (Met1, Arg6, Gln13, Gln30, Gln33, Met41, Arg64, Arg72, Ser74, Thr107, Asp112, Lys120, His137, Glu203, Ser222, Asp274, and Glu294) exhibited alternate conformations in their side chains. The overall structure of EstS1 comprises 8 β-strands (Asp48 to Thr56; Gly59 to Ile67; Leu77 to Tyr80; Val108 to Asp112; Leu148 to Asp153; Ile174 to Phe180; Ala240 to Glu246; Thr268 to Phe277) interspersed with 12 α-helices and several loops (Figure 5A). Markedly, the β-strand from Pro60 to Tyr66 extends in an anti-parallel direction to the to the other strands, similar to other esterase enzymes (Figure 5A) (18).

**Table 1:** Statistics of data refinement of native as well as complex structures of EstS1.

| Property | (native 1.22 Å) | (native 1.5 Å) | (Jeffamine) | (para-nitrophenol) | (para-nitrophenyl butyrate) | (Monomethyl phthalate) | (Dimethyl phthalate at active site) | (Dimethyl phthalate at surface) | (Diethylhexyl phthalate S154A mutant) | (Monoethylhexyl phthalate) | (Phthalate) |
| --- | --- | --- | --- | --- | --- | --- | --- | --- | --- | --- | --- |
| Space group | P 63 | P 63 | P 63 | P 63 | P 63 | P 63 | P 63 | P 63 | P 63 | P 63 | P 63 |
| Resolution (Å) | 25.8–1.2 | 25.9–1.5 | 25.8–1.5 | 23.4–1.5 | 23.4–1.8 | 25.6–2.0 | 25.5–1.6 | 25.6–1.5 | 23.1–2.4 | 25.9–1.9 | 27.1–1.5 |
| % Data completeness (in resolution range) | 100.0 (25.9–1.2) | 99.9 (25.9–1.5) | 100.0 (25.9–1.5) | 99.9 (23.4–1.5) | 99.9 (23.4–1.8) | 99.9 (25.9–2.0) | 100.0 (25.5–1.6) | 99.9 (25.6–1.5) | 99.8 (23.1–2.4) | 99.8 (25.9–1.9) | 100.0 (27.1–1.5) |
| $R_{merge}$ | 0.12 | 0.15 | 0.05 | 0.12 | 0.14 | 0.14 | 0.17 | 0.15 | 0.14 | (Not available) | 0.08 |
| $\langle I/\sigma(I) \rangle$ | 1.33 (at 1.20Å) | 1.46 (at 1.50Å) | 3.01 (at 1.50Å) | 1.68 (at 1.50Å) | 2.39 (at 1.80Å) | 5.56 (at 1.99Å) | 1.78 (at 1.60Å) | 1.82 (at 1.50Å) | 3.72 (at 2.41Å) | 1.76 (at 1.91Å) | 1.54 (at 1.50Å) |
| $R$, $R_{free}$ | 0.173, 0.204<br>0.178, 0.211 | 0.151, 0.205<br>0.162, 0.209 | 0.117, 0.163<br>0.119, 0.165 | 0.126, 0.177<br>0.127, 0.177 | 0.122, 0.185<br>0.123, 0.184 | 0.108, 0.201<br>0.110, 0.200 | 0.147, 0.194<br>0.184, 0.224 | 0.151, 0.200<br>0.152, 0.201 | 0.173, 0.246<br>0.174, 0.247 | 0.184, 0.218<br>0.193, 0.220 | 0.118, 0.163<br>0.118, 0.163 |
| $R_{free}$ test set | 4733 reflections (5.15%) | 2359 reflections (4.94%) | 2361 reflections (4.95%) | 2365 reflections (4.94%) | 1276 reflections (4.59%) | 963 reflections (4.72%) | 1848 reflections (4.73%) | 2351 reflections (4.93%) | 589 reflections (5.00%) | 1164 reflections (4.92%) | 2385 reflections (4.95%) |
| Wilson B-factor (Å<sup>2</sup>) | 6.1 | 10.1 | 11.0 | 9.9 | 15.2 | 15.1 | 12.0 | 15.0 | 21.9 | 24.2 | 8.7 |
| Anisotropy | 1.437 | 0.789 | 0.588 | 0.325 | 0.199 | 0.436 | 0.318 | 0.333 | 0.065 | 0.099 | 1.118 |
| Bulk solvent $k_{sol}(\text{e}/\text{Å}^3)$, $B_{sol}(\text{Å}^2)$ | 0.45 , 50.0 | 0.44 , 41.1 | 0.45 , 52.9 | 0.48 , 54.3 | 0.46 , 50.0 | 0.47 , 49.6 | 0.48 , 48.5 | 0.44 , 40.9 | 0.39 , 52.6 | 0.46 , 49.1 | 0.43 , 56.0 |
| $F_o, F_c$ correlation | 0.97 | 0.97 | 0.98 | 0.98 | 0.97 | 0.97 | 0.96 | 0.97 | 0.94 | 0.96 | 0.98 |
| Total number of atoms | 5128 | 4858 | 5067 | 5167 | 4865 | 4771 | 5055 | 4911 | 2440 | 4818 | 5104 |
| Average B, all atoms (Å<sup>2</sup>) | 15.0 | 17.0 | 18.0 | 16.0 | 21.0 | 20.0 | 10.0 | 20.0 | 23.0 | 30 | 16.0 |

**Figure 5:**
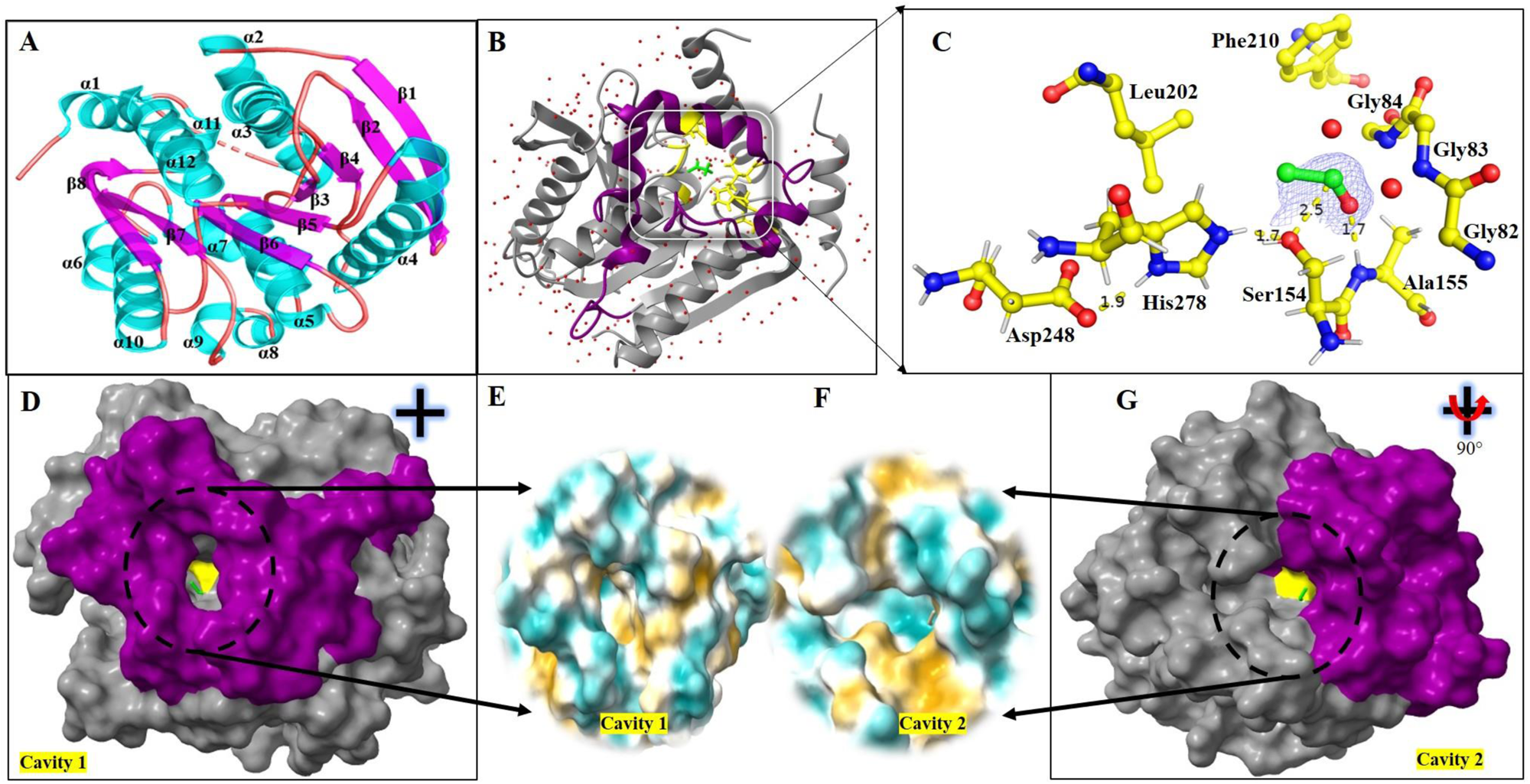
Crystal structure of EstS1 esterase at 1.22Å. A). Cartoon representation of EstS1 esterase showing α/β hydrolase fold consisting of 8 β-sheets interspersed by 12 α-helices with second β-strand running antiparallel to others. B). Cartoon representation of EstS1 esterase with active site residues (yellow) in stick representation, water molecules as red spheres, and cap domain in purple colour. C) Popout active site view in ball and stick representation of EstS1 esterase showing electron density F_o_-F_c_ map contoured at 3σ of acetyl group bound to the active site serine at a distance of 2.5Å providing a catalytically active conformation to the active site residues where the hydroxyl group of Ser154 interact with NH ɛ-atom of His278, while the O δ2 atoms of Asp248 is 1.9 Å away from the NH δ1 atom of His278. D). Surface representation of cavity 1 observed in EstS1 structure formed by the cap domain (purple) (Active site visible inside the cavity is marked in yellow color) E) Hydrophobic surface of the cavity 1 showing majority of highly hydrophobic residues (brown) inside the cavity while hydrophilic (blue) environment surrounding the cavity. F) Hydrophobic surface of cavity 2 shows a majority of hydrophobic residues (brown) on the outer side of the cavity while hydrophilic residues (blue) are present inside the cavity. G) Surface representation of cavity 2 found at the interface of cap domain and α/β hydrolase fold (Active site visible inside the cavity is marked in yellow colour).

### Cap domain covering the active site

The lid or cap domain, extending from residue Tyr181 to Pro239 consisting of 4 small α-helices connected by loops, (Figure 5B) covers the catalytic triad (Ser154-His278-Asp248) (Figure 5C) of the esterase and forms a cavity directing towards the catalytic site (Figure 5D). Two residues, Leu202 and Phe210 within the cap domain lie close to the catalytic triad and may play a role in proper orientation of substrate into the active site (Figure 5C) (18). The surface cavity 1, formed by the cap domain and leading to the active site, likely facilitates the entry of hydrophobic substrates into the active site cavity due to its hydrophobic interior (Figure 5D). This cavity extends through the catalytic site and opens up on the surface as cavity 2 at the interface of the cap domain α7 helix and the α2 helix of esterase. The arrangement of amino acids around these cavities suggests a hydrophobic environment within the cavities and the tunnel formed between them that passes through the catalytic site (Figure 5E and F). The missing loop (Pro16 to Gly22) potentially covers cavity 2, although its precise orientation could not be determined due to the absence of density in the electron density map. It is likely that the high flexibility of this loop may facilitate the exit of the product from the active site post-catalysis (Figure 5G).

### Architecture of Catalytic site

The catalytic site of EstS1 consists of the catalytic triad of EstS1, Ser154, His278, and Asp248, places the active site residues on different loop regions (Figure 5C). The catalytic Ser154 is positioned between the 5^th^ β-strand and the following α-helix, exhibiting a lower B-factor value of 8.15 Å^2^ indicating a rigid geometry similar to other α/β hydrolases. This region is known as the nucleophile elbow (35). The conserved G-X-S-X-G motif contains the catalytic Ser154 (36), while Asp248 is positioned on the loop connecting the 7^th^ β-strand to the following α-helix. His278 is located on the loop following the last β-strand. This unique arrangement of residues, despite their distant location in primary sequence and the exclusive arrangement of these distantly located residues, allows them to form the active site. Within the catalytic triad, polar contacts were observed, with the hydroxyl oxygen atom of Ser154 being 1.7 Å from the NH ɛ-atom of His278 and the O δ2 atoms of Asp248 being 1.9 Å from the NH δ1 atom of His278, indicating significant stable interactions (Figure 5C). Additionally, two water molecules near the active site Ser154 were identified, which are crucial for substrate release from the active site as described in the canonical esterase mechanism (37).

### Unexpected electron density of acetyl group

Surprisingly, electron density corresponding to an acetyl group was observed in Fourier and difference Fourier maps contoured at 3σ, located above the catalytic Ser154 at a distance of 2.50 Å (between the carbonyl carbon of the acetyl group and the hydroxyl oxygen of Ser154). An O-N bond was formed between the oxygen of the acetyl group and the NH of the Ala155 backbone at a distance of 1.7 Å (Figure 5C). This suggests the presence of the characteristic tetrahedral intermediate of the carboxylesterase mechanism (Figure 1) (37). The presence of similar extra density has been observed in the structure of Est30 carboxylesterases, suggesting the formation of a tetrahedral intermediate during the ester hydrolysis (25). Both potential tetrahedral intermediates observed in canonical esterase mechanism were analyzed, revealing that difference Fourier map and geometrical constraints align with the presence of an acetyl group (CH3-CO) with a molecular weight of 43 Da. The B-factor values for the atoms in the bound acetyl group were 13, 23, 28, and 29 Å^2^, respectively. The intermediate state is stabilized through hydrogen bonding with Ala155, which, along with Gly83 and Gly84, forms the oxyanion hole (37) (Figure 1).

Notably, no additional acyl esters were added during the process of protein expression, purification, or crystallization. Therefore, it is likely that the acetyl group was acquired from bacteria during the protein’s expression in *E. coli*.

### DEHP degradation at EstS1 active site

Co-crystallization of wildtype EstS1 with DEHP demonstrated the catalytic activity of the enzyme in its crystal form. The difference Fourier map contoured at 3σ accurately displayed DEHP degradation products monoethylhexyl phthalate and 2-ethylhexanol (Figure 6A and C). The crystal structure was solved at 1.9 Å with R*_work_* and R*_free_* values of 18.4 and 21.8%, respectively. The monoethylhexyl phthalate was located at the opening of cavity 2 extending from the catalytic site surrounded by residues Gln33, Leu37, Gln40, and Met41 (Figure 6B, E and F). Meanwhile, 2-ethylhexanol was observed near the opening of cavity 1, leading to the catalytic or active site (Figure 6E and G). The residues observed in the close vicinity of 2-ethylhexanol were Met207, Phe210, Gly211, and Lys193 (Figure 6D). Similarly, electron density of product has been observed in previous studies involving co-crystallization with substrates or inhibitors (8, 38).

**Figure 6:**
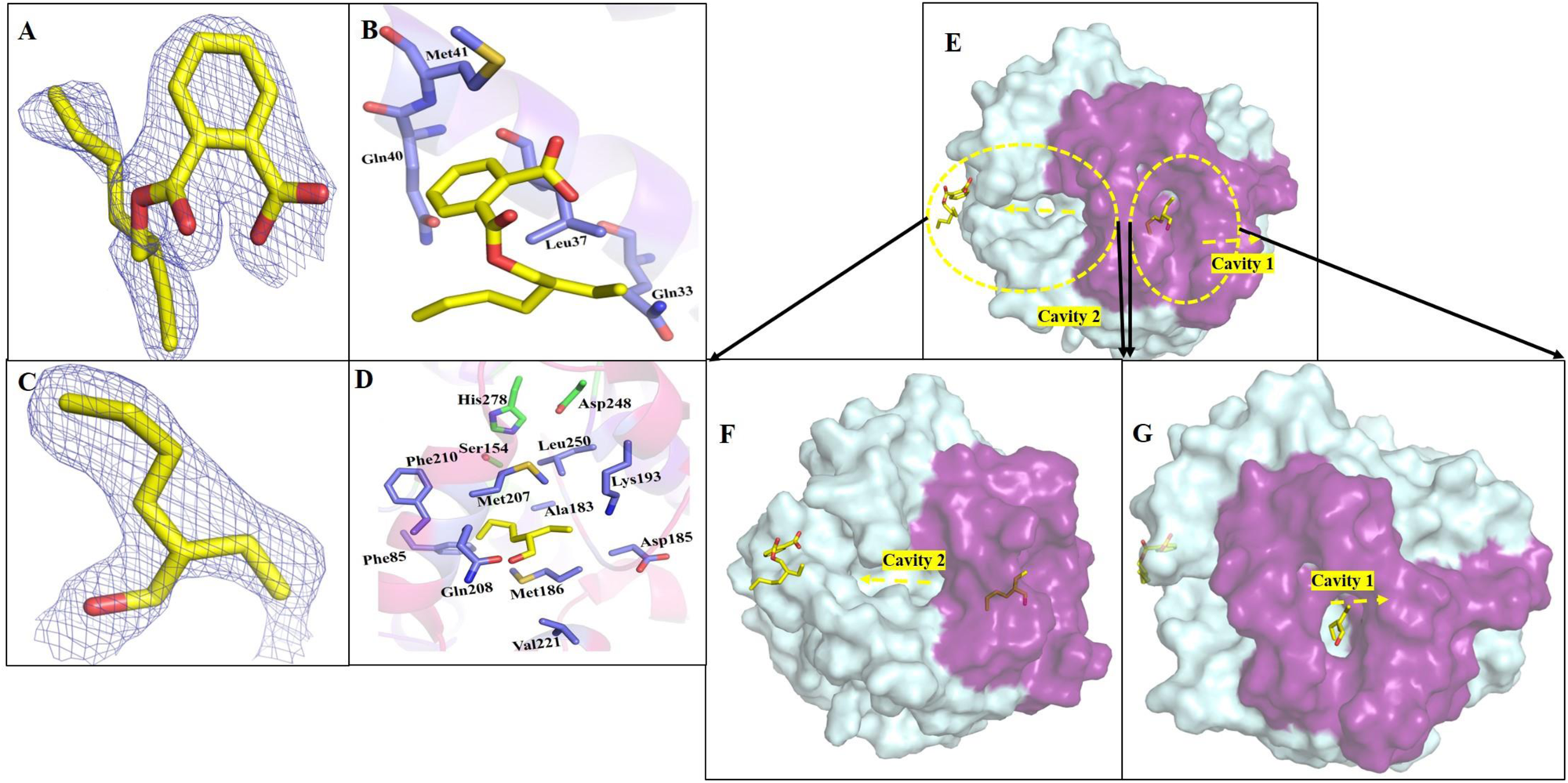
Crystal structure of DEHP degradation products MEHP and 2-ethyl hexanol with wildtype EstsS1 esterase at 1.9Å. A) Electron density F_o_-F_c_ map contoured at 3σ of MEHP (yellow) B). Stick representation revealing interactions site of MEHP after leaving the catalytic tunnel. C) Electron density F_o_-F_c_ map contoured at 3σ of 2-ethyl hexanol (yellow). D) Stick representation showing the interaction site of 2-ethyl hexanol after leaving the catalytic tunnel. E) Surface representation of EstS1 in complex with MEHP and 2-ethyl hexanol displaying the relative position of the two with respect to the cavity 1 and 2 of the catalytic tunnel. The MEHP was observed to be bound at the outer surface of cavity 2 after leaving the catalytic site while 2 ethyl hexanol was observed to be bound at the outer surface of cavity 1. F) Popout view showing the position of MEHP while coming out of the cavity 2 G) Popout view showing the position of 2-ethyl hexanol while coming out of cavity 1. (Yellow dashed arrows represent the direction of entry and exit of the molecule, and purple colour surface denotes the cap domain).

### DEHP in complex with Ser154Ala mutant

To obtain a complex structure of DEHP with EstS1, a Ser154Ala substitution mutation was introduced at the key catalytic residue. The co-crystallized structure of DEHP with Ser154Ala mutant of EstS1 was solved at 2.4 Å, with R*_work_* and R*_free_* values of 17.3% and 24.6% (Figure 7A). The orientation of DEHP at the active site was similar to the tetrahedral complex of the canonical esterase mechanism. The carbonyl oxygen of the ester group formed a hydrogen bond (2.8 Å) with the backbone nitrogen of the Gly83 residue, which forms the oxyanion hole (Figure 7 B and C). The ester oxygen formed a hydrogen bond (3.4Å) with the ɛN of His278 due to the absence of the OH group of the catalytic serine (Figure 7B). Additionally, three water molecules, shown as red spheres in Figure 6C, formed several hydrogen bonds with carbonyl and ester oxygen atoms of DEHP, as well as with other residues in the binding cavity, including Ala155, Gly84, and Gly83 of the oxyanion hole. This arrangement of water molecules in the active site cavity stabilizes the binding of DEHP at the catalytic site and suggests a water-bound acyl state during esterase catalysis (Figures 6D, E, and F).

**Figure 7:**
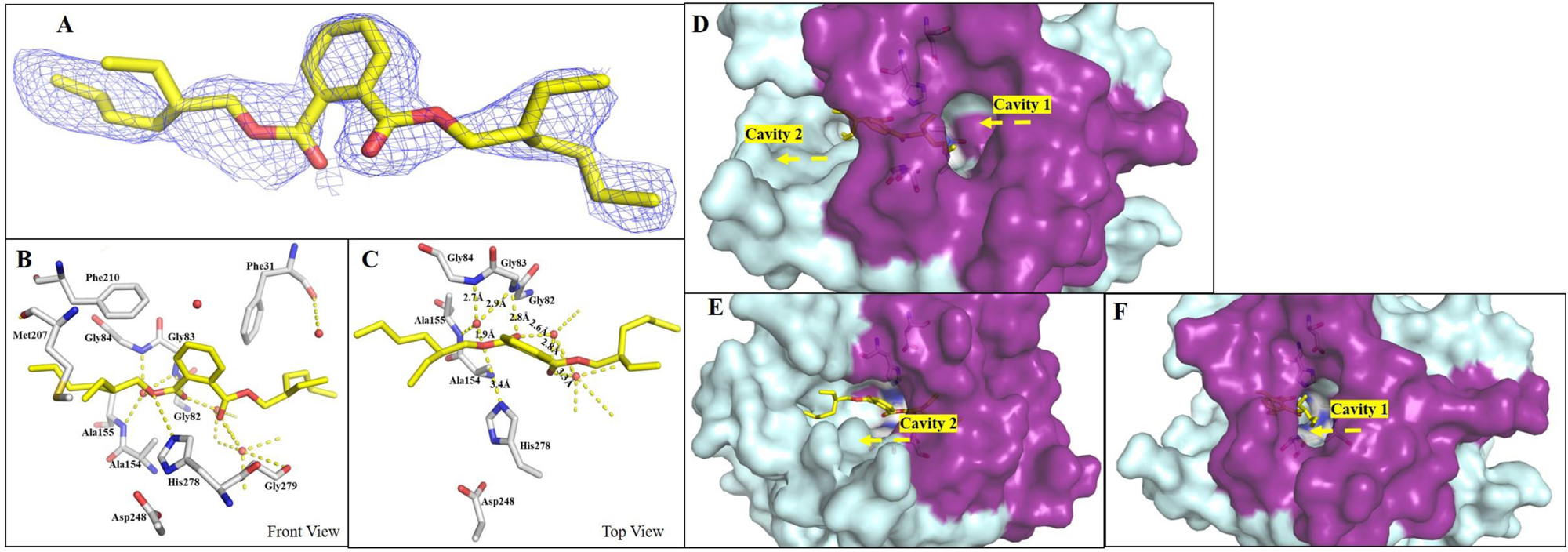
Crystal structure of DEHP-EstS1 S154A mutant complex at 2.4Å showing the effective conformation of DEHP at the active site of EstS1. A) Electron density F_o_-F_c_ map contoured at 3σ of DEHP at the active site of EstS1. B). Stick representation (front pose) of DEHP (yellow) at active site showing interactions with the catalytic residues (grey) of EstS1 and water molecules (red spheres) (yellow dash line marks the bond and bond distances) C) Stick representation (behind pose) of DEHP (yellow) at active site showing interactions with the catalytic residues (grey) of EstS1 and water molecules (red spheres) (yellow dash line marks the bond and bond distances) D) Surface view of DEHP-EstS1 S154A mutant complex showing DEHP orientation in the catalytic tunnels between cavity 1 and cavity 2. E) Surface view of DEHP-EstS1 S154A mutant complex showing the orientation of DEHP with respect to cavity 2. F) Surface view of DEHP-EstS1 S154A mutant complex showing DEHP orientation with respect to cavity 1. (Yellow dashed arrows represent the direction of entry and exit of the molecule, red sphere represents water molecules and purple color surface denotes the cap domain).

### Tracing the active site tunnel architecture from entry to exit

To thoroughly investigate the structural features associated with the substrate’s pathway to the active site and release of its product from the active site tunnel, which lies between two surface cavities on EstS1, multiple crystal structures of EstS1 complexed with various substrates, products, and their analogs were determined (Figure S4, S5 and 8). High-resolution structures of EstS1 in complex with dimethyl phthalate (1.6 Å and 1.5 Å) (Figure S4 A-F and Figure S4 G-I), para-nitrophenyl butyrate (1.8 Å) (Figure S4 J-N), monomethyl phthalate (2 Å) (Figure S4 O-T), para-nitrophenol (1.5 Å) (Figure S5 A-D), phthalate (1.5 Å) (Figure S5 E-G), and jeffamine (1.5 Å) (Figure S5 H-K) revealed multiple densities of these ligands within the active site tunnel, providing detailed evidence of their pathways within EstS1 esterase.

The co-crystallized structure of EstS1 with dimethyl phthalate, solved at 1.6 Å resolution (R*_work_* 14.7%, R*_free_* 19.4%), showed difference Fourier maps contoured at 3σ of dimethyl phthalate at two sites: one near the active site residues (Figure S4 A) another near the surface at cavity 2 (Figure S4 D). Key residues near the second-density map include Leu37, Gln40, and Met41 (Figures S4 D and E). Another structure at 1.5 Å revealed dimethyl phthalate bound at the enzyme’s outer surface near a small cleft formed by Arg216, His223, Pro122, and Trp225, which contribute to the enzyme’s cap domain (Figure S4 G and H). This cleft leads towards the active site cavity (Figure S4 I). An additional difference Fourier map for the acetyl group was observed at the active site in both structures, interacting with the catalytic Ser154 and oxyanion hole residues, resulting in hindrance in catalytic degradation of dimethyl phthalate at the catalytic Ser154 residue.

The EstS1-p-nitrophenyl butyrate complex structure, solved at 1.8 Å resolution (R*_work_* 12.2%, R*_free_* 18.5%), revealed difference Fourier maps contoured at 3σ of two pNPB molecules. One molecule was on its way to reach the active site, with para-nitro group away from the active site residues and the butyrate end closer to Ser154, with the C1 carbon of pNPB 2.4 Å from the OH group of catalytic serine and the ester oxygen 4.5 Å away (Figure S4 J and K). The other molecule was further along, interacting with residues in the cap region near cavity 1 (Figure S4 K, L, M, and N). The yellow coloration in the crystal well solution indicated the conversion of pNPB (colorless) to pNP (yellow) but the density map for the product was not observed in the crystal structure due to the high pace of conversion and release of product from the catalytic site.

The EstS1-monomethyl phthalate complex structure, solved at 2 Å (Rwork 18.4%, Rfree 21.8%), showed difference Fourier maps contoured at 3σ of monomethyl phthalate at three positions (Figure S4 O and R). One position was in between the active site and cavity 1, suggesting ligand entry via surface cavity 1 (Figure S4 P and T). The second position of monomethyl phthalate was observed at the active site, forming hydrogen bonds between its ester oxygen and His278 at 2.0 Å (Figure S4 P and Q). The carbonyl oxygen of monomethyl phthalate formed hydrogen bonds with Gly83, Gly84, and Ala155 residues forming the oxyanion hole (Figure S4 P). The third density map was observed in a small cleft involving Lys32, Asp51, Arg62, Ile63, Arg64, Asp89, Ile90, Asp91, and Asp112 (Figure S4 R). These conformations illustrated the ligand’s pathway through the catalytic tunnel, highlighting the cap domain’s role in substrate specificity (Figure S4 Q and T).

The EstS1-pNP complex, determined at 1.5 Å, revealed difference Fourier maps of pNP contoured at 3σ at two positions (Figure S5 A). One was in a cleft between alpha helices of the cap region (Leu215 to Pro218), with nearby residues including Gln208, Gly211, Glu212, Leu215, Arg216, Thr217, and Pro218 (Figure S5 B). The second was at the active site, with the hydroxyl group of pNP’s phenol ring forming a bond with Ser154 at 2.6 Å (Figure S5 B). The positioning suggested substrate entry through cavity 1 (Figures S5 C and D). The acetyl group was observed near Ser154 forming hydrogen bonds with Ala155, Gly83, and Gly84, mimicking the tetrahedral conformation of the acyl-enzyme intermediate (Figure S5 B).

In the EstS1-phthalate complex, solved at 1.5 Å, phthalate was observed in two positions within the catalytic tunnel (Figure S5 E). One orientation was bound at the active site, forming hydrogen bonds between phthalate’s O8 oxygen and His278’s ɛN at 2.7 Å, and the O9 oxygen formed hydrogen bonds with Gly83, Gly84, and Ala155, resembling the typical esterase reaction intermediate (Figure S5 F). Water molecules contributed to the stability of this orientation, suggesting cavity 2 as a potential exit pathway for the product post-catalysis. The other density was near cavity 1, indicating the entry pathway (Figure S5 G).

The EstS1-jeffamine complex structure, solved at 1.5 Å, showed jeffamine entering the active site through a loop between cap region helices (Figure S5 H). Jeffamine interacted with Glu212-Pro218 residues and Ser154, forming a covalent bond (1.7 Å) with the hydroxyl oxygen of serine and the terminal oxygen of the propylene monomer (Figure S5 I). Jeffamine’s oxygen also formed a hydrogen bond with His278 at 3.2 Å, indicating the interaction potential of EstS1 with polypropylene (Figure S5 J and K).

The EstS1-jeffamine complex structure, solved at 1.5 Å, showed Jeffamine, a long polymer of polypropylene glycol, entering the active site through a loop between cap region helices determining the entry site of catalytic tunnel (Figure S5 H). The outer end of jeffamine interacted primarily with the Glu212-Pro218 residues while the other end interacted with the active site Ser154, forming a covalent bond at a distance of 1.7 Å between the hydroxyl oxygen of serine and the terminal oxygen of the propylene monomer of jeffamine (Figure S5 I). The same oxygen atom of jeffamine that interacted with serine also formed a hydrogen bond with the His278 at a distance of 3.2 Å. This complex structure of jeffamine with EstS1 displays effective interaction with the active site residues, indicating the interaction potential of the enzyme with polypropylene to mediate its degradation (Figure S5 J and K). Therefore, the complex of EstS1 with jeffamine not only presented the path of followed by the substrate towards the active site but also revealed the broad substrate specificity of EstS1.

These crystal structures provided comprehensive insights into substrate entry, catalysis, and product exit from the EstS1 active site cavity. A comparative analysis suggested cavity 1 as the entry site and cavity 2 as the exit site for substrates and products, respectively (Figure 8). Superimposition of active site residues across different structures showed a rigid conformation of the catalytic triad, while a significant angular movement (134°) was observed in Met207’s side chain in some complexes. This movement was seen in complexes with both substrates and products, making it difficult to conclude its role in regulating catalysis (Figure 8C). Further mutational analysis of this residue was suggested. Additionally, a minor angular deviation (16°) was noted in Phe31, which is distant from the catalytic triad.

**Figure 8:**
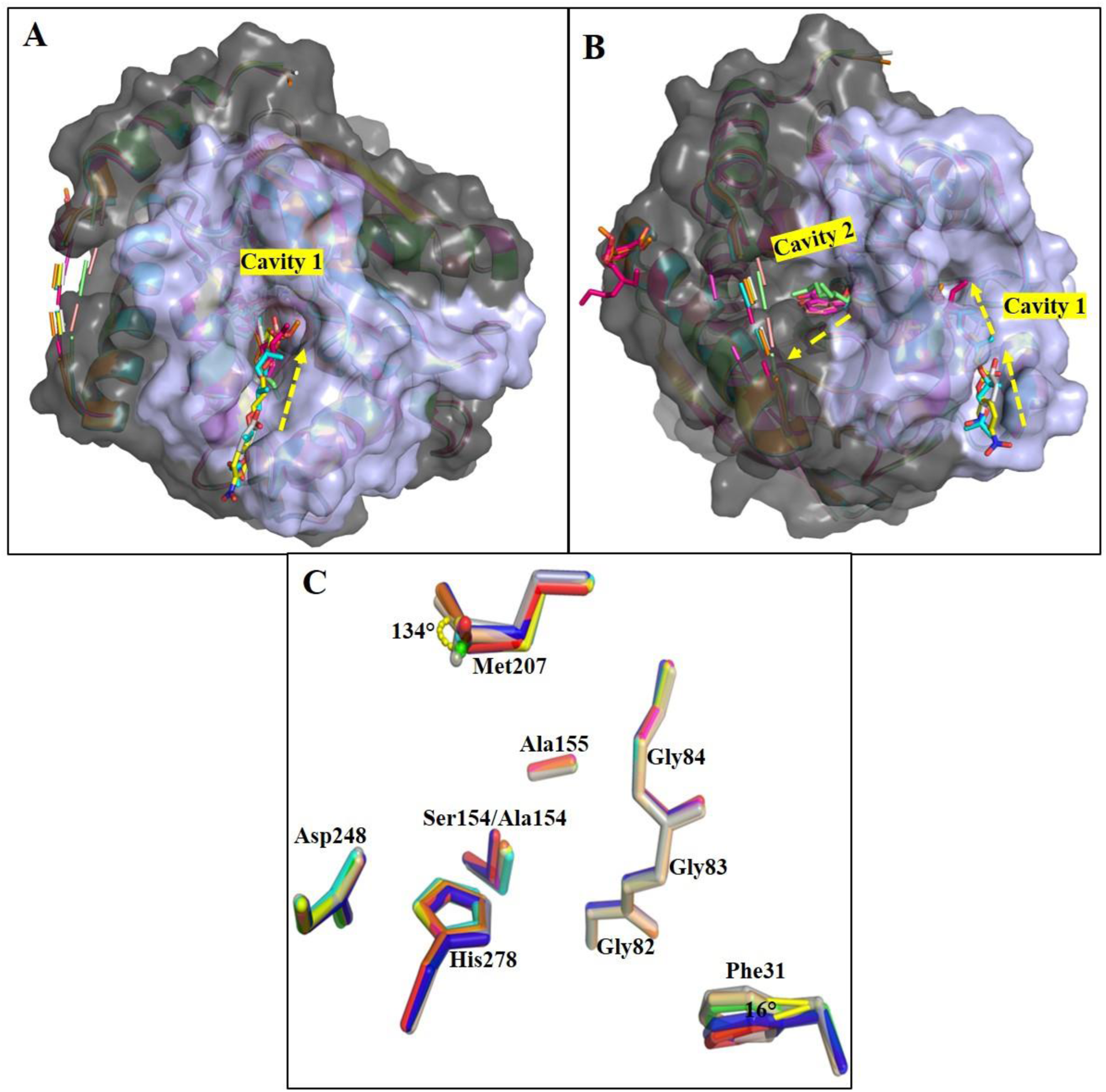
Superimposition of all the EstS1 complex structures over EstS1 apo structure displaying structural changes due to the presence of ligand molecules bound to it. A) Surface representation of superimposed complex structures showing the entry site of all the ligands through cavity 1 formed by the cap domain (blue). B) Surface view of the entrance cavity 1 displaying the position of all the ligands entering into the catalytic tunnel and exit cavity of EstS1 showing the position of monoethylhexyl phthalate (orange) at the outer side of the exit cavity and location of other ligands into the catalytic tunnel moving towards the exit site C) Relative conformation of active site residues in native (cyan) as well as complex structures (EstS1-DEHP: grey; EstS1-monoethylhexyl phthalate complex: wheat; EstS1-Dimethyl phthalate at active site: orange; EstS1-Dimethyl phthalate on surface: blue; EstS1-pNPB:green; EstS1-monomethyl phthalate: dark blue; EstS1-pNP: mauve; EstS1-phthalate: salmon red; EstS1-jeffamine: yellow) displaying angular displacement in binding site residues Met207 and Phe31

### Methionine 207: A hidden player in EstS1 catalysis

Based on the detailed structural analysis, site-directed mutagenesis was conducted to investigate the roles of binding site residues in EstS1 catalytic activity and to identify substitutions that could enhance its catalysis. Two mutants, Ser154Thr and Met207Ala, were created to assess their effects on DEHP binding at the EstS1 substrate binding site. Substituting the active site residue Ser154 with threonine reduced DEHP binding, as evidenced by decreased fluorescence quenching in the mutant compared to the wild type (Figure 2B). Surprisingly, replacing Met207 with alanine, a mutation inspired by the observed positioning of Met207 relative to the substrate, resulted in almost no DEHP binding at the active site, as indicated by minimal fluorescence quenching (Figure 2B). This result highlighted Met207’s crucial role in facilitating DEHP binding at the EstS1 active site, establishing it as an important residue for catalysis.

This significant finding from mutational analysis provides valuable insights that could be useful for further enzyme engineering studies.

### Molecular dynamic simulation of the EstS1-DEHP complex

To study the dynamics of the interactions between DEHP and wildtype EstS1, and to assess the binding affinity and stability of these interactions, molecular dynamics (MD) simulations were conducted for both the apo-EstS1 and the EstS1-DEHP complex crystal structures for 100 ns. The missing loop region in the protein structure was modeled using Coot software, and the alanine residue present in the Ser154Ala mutant crystal structure of EstS1-DEHP complex was replaced with the catalytic serine. Analysis of the MD simulation trajectory showed minimal average fluctuations in the protein residues in the presence of DEHP compared to apo-EstS1, as indicated by the Root Mean Square Fluctuations (RMSF) versus residue plot (Figure S6 A). The Root Mean Square Deviations (RMSD) values over time varied within a narrow range of 0.1 to 0.275 nm, indicating highly stable interactions of DEHP within the enzyme’s active site cavity (Figure S5B). The solvent-accessible surface area (SASA) slightly increased in the presence of DEHP compared to apo-EstS1 (Figure S6 C), and a slight increase in the radius of gyration (Rg) values was observed, suggesting a minor decrease in the protein’s compactness when DEHP was bound (Figure S5D) (39).

Secondary structure analysis of both the DEHP complex and the apo-protein showed minimal changes in the secondary structure in the presence of DEHP, indicating stable binding of DEHP to EstS1 and a rigid geometry of the active site tunnel (Figure S6 E and F). The free energy landscapes of apo-EstS1 and the DEHP-EstS1 complex revealed that DEHP binding steepened the landscape and reduced free energy barriers, stabilizing the protein in a more native folded state compared to the high fluctuations and multiple local minima observed in the apo-protein, which correspond to transition or metastable states close to the native folded state (Figure S7 A and B) (40).

The MM/PBSA free binding energy analysis of the last 40 ns of the simulation trajectory demonstrated stable interaction of DEHP with EstS1, with a total negative binding energy ranging from −20 to −45 kcal/mol (Figure S7 C). Cluster analysis of the trajectory showed effective clustering, with a large number of conformations in the top 10 clusters. Superimposition of the top 10 cluster conformations of the DEHP-EstS1 complex revealed strong interactions of DEHP with the active site residues and those forming the oxyanion hole, illustrating the typical esterase mechanism (Figure S7 D-F).

Quantum Mechanics/Molecular Mechanics (QM/MM) GBSA analysis, including residues of the catalytic site (Gly82, Gly83, Gly84, Ser154, Ala155, Asp248, and His278) and DEHP in the quantum mechanical region, confirmed the stable interaction and binding of DEHP at the catalytic site of EstS1 esterase, with a net negative total energy component ranging between - 25 to −42 kcal/mol, mainly contributed by van der Waals interactions (Figure S8) (39).

## Discussion

Phthalate esters, including bis(2-ethylhexyl) phthalate (DEHP), are widely used in personal care products and are persistent environmental contaminants with known adverse effects on human health and ecosystems (1). Several enzymes capable of degrading small molecular weight phthalate diesters have been reported, but there is limited knowledge about enzymes that can effectively degrade high molecular weight phthalate diesters (7, 10, 13, 14). EstS1, a thermostable and pH-tolerant esterase, is known for degrading low molecular-weight phthalate diesters, but its ability to degrade high molecular-weight phthalates has not been established (13). Moreover, to date, structural information about high molecular weight phthalate diester degrading esterases is scanty. This study aims to fill these gaps by providing a biochemical, biophysical, and structural characterization of the dialkyl phthalates-degrading esterase EstS1, particularly its potential to degrade DEHP, a high molecular weight phthalate diester.

Biophysical and biochemical analysis disclosed significant binding and degradation of DEHP by EstS1, achieving a degradation rate of 390 g/L in 2 days, which is comparable to previous studies (12, 7). The high-resolution crystal structures of EstS1 in both native and complex forms lay the foundation for understanding the catalytic mechanism involved in the degradation of a wide range of phthalate diesters. EstS1 has the ability to accommodate both aliphatic and aromatic esters indicating its broad substrate specificity, making it valuable for degrading various plasticizers under in-situ conditions. Mutational analysis of the active site could potentially alter substrate specificity, further expanding the enzyme’s substrate range (41).

### EstS1 catalytic architecture and substrate specificity

Structural analysis revealed conserved catalytic site residues forming a catalytic triad (Ser154-His278-Asp248), suggesting its resemblance with other carboxyl esterases (13). However, significant variations were observed in the cap domain, which is likely responsible for the enzyme’s substrate specificity (22). In comparison with MEHP hydrolase (8), the length of the cap domain of EstS1 covering the active site is less, which in turn expands the exposure of the active site’s surface to substrates. The cap domain (Tyr181-Pro239) (Figure 5), enables EstS1 to interact with both aromatic and aliphatic compounds, showcasing its broad substrate specificity.

Further examination of EstS1 for structural homology within the hormone-sensitive lipase (HSL) family highlighted the diversity of cap domains in this enzyme family. Superimposing the structure of EstS1 with PestE (43), another HSL-like family esterase, revealed key alpha-helical conformation similarities. Additionally, unmodeled electron density near Ser154, similar to observations in PestE (43), hinted at a tetrahedral intermediate involving acyl binding. This phenomenon highlights the significance of the observed extra density at the active site of EstS1, suggesting the presence of an acetyl group, which could be attributed to the formation of a tetrahedral complex (25, 43).

### DEHP degradation mechanism, and mutation analysis

Notably, the soaking of EstS1 S154A mutant crystals with DEHP for 30 minutes resulted in the difference Fourier map of DEHP contoured at 3σ observed at the active site. In contrast, co-crystallization of DEHP with the wildtype EstS1 for 2 days revealed densities in difference Fourier map corresponding to the degradation products, monoethyl hexyl phthalate and 2-ethylhexanol, at the active site tunnel’s exit and entry sites, respectively. This provides clear evidence for the DEHP degradation potential of EstS1.

Esterases generally follow a bi-bi ping-pong mechanism involving an acyl-serine intermediate, characterized by a catalytic triad, an oxyanion hole, and a specific lid or cap domain. The oxyanion hole stabilizes the tetrahedral intermediate through hydrogen bonds between the substrate’s carbonyl oxygen and two main-chain N-H groups. The third hydrogen bond donor varies among different enzymes. The enzyme is acylated via a first tetrahedral intermediate, followed by the release of the acyl group through a second intermediate involving a water molecule (18, 20).

The EstS1 Ser154Ala-DEHP complex’s carbonyl oxygen formed a hydrogen bond with Gly83, and a nearby water molecule stabilized the complex. However, due to the absence of active site Ser154, the exact tetrahedral complex was not observed. The catalytic triad maintained a rigid arrangement across all complex structures, consistent with minimal reorganization during catalysis (18, 44). Additionally, the Met207 residue, located just above DEHP in the complex structure, exhibited significant deviations, along with Phe31, which showed mild deviations in some complexes. Furthermore, the biophysical experiment showed a reduction in the binding of DEHP when Met207 was mutated to alanine thus disclosing its role in enhancing the binding of DEHP by providing it with proper orientation for the catalysis (45).

### Tracing the path from entry to exit

The presence of multiple difference Fourier map densities corresponding to substrates and products has unveiled critical insights into the trajectory followed by these molecules as they approach and depart from the active site. The complex structures of EstS1 with various substrates and products provided insights into the catalytic mechanism and substrate/product entry and exit pathways. An open cavity formed by alpha-helical structures of the cap domain interspersed by loop residues mediates the entry of substrate into the catalytic tunnel and thus was identified as the entry site (Cavity 1) for substrates. An exit site (Cavity 2) for products was revealed near Leu37, Glu40, and Met41. A corresponding density of the by-product has been observed in monoalkyl phthalate esterase suggesting a similar exit tunnel for release of product from the catalytic site (8). The complex of jeffamine, a polypropylene analog, with EstS1 traced the catalytic tunnel and represented the exact route followed by the substrate toward the active site. Interestingly, jeffamine was observed to form a covalent linkage with the catalytic Ser154, since it’s an analog of polypropylene the binding of jeffamine highlights the broader specificity of the enzyme. This is the first time an analog of propylene has been found to bind with esterase thus provided scope for further application of EstS1 for the degradation of polypropylene-derived polymers which could be further investigated.

### Summary

This study presents EstS1 as an efficient enzyme for degrading high molecular weight phthalate esters, specifically DEHP, demonstrating remarkable catalytic efficiency. The high-resolution crystal structures of EstS1 in both native and complex forms with various substrates, including DEHP, and products elucidate its catalytic mechanism. The complex structures, showing multiple densities for the bound substrates and products, provide detailed insights into the pathway followed by the substrate to reach the catalytic site and the exit route of the product after catalysis. Noteworthily, the structural analysis of the EstS1-jeffamine complex suggests EstS1 as a promising candidate for investigating polypropylene degradation.

The biochemical and structural studies revealed novel features of EstS1, including broad substrate specificity, a unique catalytic site architecture, and unprecedented insights into reaction intermediates. These findings collectively establish EstS1 as a promising candidate for bioremediation and industrial applications. The structural information provided in this study lays the groundwork for future AI-based computational studies and enzyme engineering efforts aimed at developing innovative solutions to address environmental challenges posed by phthalate esters.

## Materials and Methods

### Chemicals, reagents, and bacterial strains

All reagents and chemicals used for experiments were of analytical grade. Purified water for buffers with a minimum resistance of 18.2 MΩ was generated using the Merck Synergy Water Purification System. The recombinant plasmid integrated with wild-type EstS1 gene sequence and the primer sequences for site-directed mutagenesis were synthesized by Gene to protein Private Limited (Lucknow, India). The *E. coli* DH5α and BL-21 strains were procured from the Microbial Type Culture Collection (MTCC) Chandigarh, India.

### Cloning, expression and purification of EstS1 and its mutants

The recombinant pet28a-EstS1 plasmids were transformed and expressed in *E coli* BL21 λ(DE3) cells in a culture of 1 L LB (Luria-Bertani) liquid medium using 0.5 mM isopropyl β-D-thiogalactopyranoside (IPTG) as inducer at optical density of 0.6 at OD_600_. The induced culture was incubated at 16 °C for 16 h (46). The cell mass was collected and suspended in 10 ml purification buffer (0.1 M phosphate buffer at pH 7.4, 10% glycerol, and 2 mM DTT). The lysis of collected cell mass was done using a Constant Cell Disruptor at a pressure of 20 psi. and the obtained cell lysate was centrifuged at 12000 rpm for 45 min. The supernatant containing the desired enzyme was then collected and incubated with a Ni-NTA affinity column pre-equilibrated with the purification buffer. Imidazole gradient ranging from 20 to 500 mM was applied to purify the protein and fractions were subsequently analyzed on an SDS-PAGE gel. (15, 16). The protein concentration was estimated using the Pierce Micro BCA protein assay kit with bovine serum albumin (BSA) as the standard.

To prepare EstS1 mutants (S154A, S154T, and M207A) the pet28a-EstS1 plasmids were amplified using respective forward and reverse primers for mutants (Table S1). The amplified product was digested using DpnI enzyme and the digested product was transformed into E. coli. DH5α cells (47). The mutant plasmids were isolated and confirmed by sequencing. The confirmed plasmids were transformed and expressed into *E coli* BL21 λ(DE3) cells and purification of the mutant proteins was performed using the above-mentioned procedure.

### Biochemical and Biophysical analysis

#### UV-spectroscopy-based analysis of EstS1 esterase activity and kinetics

The spectrophotometric biochemical activity of EstS1 was evaluated by estimating the conversion of para-nitrophenyl butyrate (pNPB) substrate into para-nitro phenol. The quantification of the product formed was done at 405 nm. The reaction mixture consisted of 30 μM EstS1 enzyme, 0.1 M phosphate buffer at pH 7.0, and varying concentrations of substrate pNPB to a total volume of 100 μL. The kinetic parameters of EstS1 were analyzed by estimating the concentration of para-nitro phenol produced at room temperature (25 °C). The enzyme’s kinetic parameters, including Vmax (maximum velocity of reaction), Km (the Michaelis-Menten constant), and k_cat_/K_m_ were determined using the Michaelis–Menten equation (15, 16). Each experiment was done in triplicates, and average values were plotted.

#### Fluorescence spectroscopy-based binding assay of EstS1 wild-type and mutants

Fluorescence spectroscopy was performed to investigate the binding of DEHP to wild-type and mutant forms of EstS1 utilizing the HORIBA FluoroMax Spectrofluorometer and quartz cuvette of 1.0 cm. Initially, to assess fluorescence quenching by DEHP binding to EstS1 wild-type as well as mutants, a 1ml reaction mixture containing 1 μM of EstS1 enzyme and 1mM DEHP was analyzed. The emission spectra were recorded using 5 nm as slit width size, excitation wavelength of 280 nm (due to the presence of tryptophan, tyrosine, and phenylalanine in the binding cavity of the enzyme), and emission spectra range of 295 to 500 nm. Eventually, to study the change in fluorescence intensity or substrate binding with increasing DEHP concentrations, different concentrations ranging from were added to the cuvette containing 2 μM EstS1(30).

#### LC-MS based analysis of bis-(2-ethyl hexyl) phthalate (DEHP) degradation

The degradation of DEHP was analysed using liquid chromatography-mass spectrometry (LC-MS). Enzymatic reactions with optimal concentrations of EstS1 enzyme and DEHP were conducted at 37 °C and for 3 h in triplicates. The reactions were terminated by altering pH to 2.5 with HCL, and the product was extracted using an equal volume of organic solvent. The organic fraction was evaporated to dryness, then resuspended in 1 ml methanol and filtered using 0.2 μm nylon filters (48). Mass Spectrometry was performed using the Xevo G2-XS QTof Quadrupole Time-of-Flight Mass Spectrometry system (Xevo G2-XS QTof Mass Spectrometer, Waters) in positive ionization mode. Chromatographic separation was done using a reverse phase C18 chromatographic column (1.7 µm, 2.1 mm X 100 mm; Acquity UPLC BEH column) with a mobile phase consisting of formic acid (0.1% v/v) in water and methanol and a flow rate of 0.3 ml/min. The injection volume was 10 μL.

For the acquisition of MS, the capillary and cone voltages were kept at 3 kV and 40 V, respectively. The cone and desolvation gas flow rates were 50 and 900 L/h. Electrospray ionization (ESI) was used, and the scan range was set to 100-1000 m/z with a scan time of 1 s.

Data processing was performed using Waters UNIFI software, utilizing a target by mass procedure. Parameters such as target match tolerance, fragment match tolerance ion ratio tolerance, and max number of targets were set to specific values. Additionally, thresholds for relative intensity, candidates per sample, and peaks per channel were defined. The expected m/z ratios for fragments of DEHP, monoethylhexyl phthalate, and 2-ethylhexanol were determined using MestReNova software and used for library preparation.

#### Quantitative and kinetic analysis of bis-(2-ethyl hexyl) phthalate (DEHP) degradation

Degradation kinetics of DEHP was studied using high-performance liquid chromatography (HPLC). The enzyme reactions (1 ml) of DEHP containing 10 μL of the EstS1 enzyme and DEHP at different concentrations were set up and incubated at 37 °C. The reactions were terminated after 3 h and extraction was done. The mobile phase consisted of methanol (85% v/v) and water (15% v/v) and the flow rate used was 0.8 ml/min. The monitoring of eluent was done by UV absorbance at 254 nm. The extracts were then analysed by Waters HPLC system consisting of photodiode array (model PDA 2998; Waters) detector and binary pump (model 1525). The separation was achieved using a C18 XBridge (Waters) column of dimension 5 μm, × 150 × 4.6 mm. Control samples without enzyme were also analysed for generating a standard curve. All reactions were set up in triplicates, and average values were used for calculations. The enzyme kinetic parameters and k_cat_/K_m_ (catalytic efficiency) were determined (14).

#### Crystallization and co-crystallization of EstS1

The EstS1 protein was concentrated to a level of 10 mg/ml and used for crystallization experiments using the sitting drop method at a temperature of 20 °C (15, 16, 47). Initial crystals were observed in the screen and the optimal crystallization condition for obtaining high-quality diffraction data was found to be a solution containing 0.1 M HEPES (pH 7.0), and 5% v/v Jeffamine ED-2001, with a protein-to-reservoir ratio of 1:1. Further, the crystals were cryoprotected in a solution containing mother liquor solution followed by flash-freezing in a steam of liquid nitrogen at a temperature of 100 K. To investigate the binding of Est1 with various compounds, crystals were soaked or co-crystallized with di-ethylhexyl phthalate, dimethyl phthalate, para-nitro phenyl butyrate, monomethyl phthalate, para-nitro phenol, and phthalate.

To further enhance the resolution of di-ethylhexyl phthalate complex with EstS1, co-crystallization of di-ethylhexyl phthalate with S154A, S154T, and M207A (binding site residue) mutant of EstS1 were performed. Screening with different crystallization conditions resulted in best hits with EstS1 S154A mutant in solution containing Tris buffer (pH 8.5) and 1.26 M ammonium sulfate. Crystallization trials for the other two mutants failed to form crystal complexes.

#### Data collection and structure solving

The data collection was done at home source using Rigaku Micromax-007 HF high intensity microfocus rotating anode X-Ray generator integrated with Rigaku Hypix 6000C detector installed at Macromolecular Crystallography Unit, Institute Instrumentation Centre, IIT Roorkee. The structures of EstS1 in native as well as complex forms were solved using CCP4i2 (49) suit and Phenix (50). Homologous structure of *Pyrobaculum calidifontis* esterase (PDB ID 2YH2), having 43% sequence identity, was used as a search model for performing molecular replacement using MOLREP (51) and Phaser MR (52) programs. Iterative model building rounds were performed using COOT (53), while the refinement was done using REFMAC (54) and phenix.refine (50). Omit maps of the structures were generated using Phenix (50). The surface and interaction images were prepared using Pymol (55) and Chimera (56).

#### Molecular dynamic simulation and molecular mechanics and quantum mechanics binding energy calculations

The crystal structure of DEHP-EstS1 S154A mutant at 2.4Å resolution was considered for molecular dynamic simulation. The missing residues were added and the active site alanine was mutated to the wild-type serine residue using Coot (53) (https://www2.mrc-lmb.cam.ac.uk/personal/pemsley/coot/) software to evaluate the actual interaction mechanism of EstS1 with DEHP. The so-formed wild-type EstS1 apo form and EstS1-DEHP complex were utilised for molecular dynamic simulation of 100 ns using GROMACS (57) (https://www.gromacs.org/) software. The topologies files of ligand (DEHP) and protein were generated by using CGenFF web server (58) (https://cgenff.com/) and CHARMM36 force field respectively. Both apo-EstS1 and EstS1-DEHP complex were solvated in a dodecahedron box. Sodium ions were added to neutralize the system. Energy minimization limited with a maximum force of 10kJ/mol was done. Equilibrations at Normal Volume Temperature (NVT) and Normal Pressure Temperature (NPT) were performed for a period of 100ps. Finally, the equilibrated systems were subjected to production MD runs of 100 ns. The MD simulation trajectory was analyzed based on variations in Root Mean Square Deviation, Root Mean Square fluctuation, Radius of Gyration, Solvent Accessible Surface Area, and changes in secondary structure. The PCA analysis was performed and the free energy landscapes of the protein in apo and complex forms were generated. The cluster analysis was performed to analyze the most frequent interactions of the ligand with the protein and snapshots of the top 10 clusters were collected and superimposed to study the most frequent structural changes and the mechanism of interaction of the DEHP with EstS1. The free binding energy analysis was performed using gmx-mmpbsa tool (59) (https://valdes-tresanco-ms.github.io/gmx_MMPBSA/dev/). To further study the interaction of the lead active site residues and their binding potentials quantum mechanics/molecular mechanism generalized Born surface area (QM/MM GBSA) free energy analysis was performed by selecting Gly82, Gly83, Gly84, Ser154, Asp248, His278 residues and DEHP in the quantum mechanics layer. The analysis of free binding energies was done using gmx_analyser. All plots were generated using Origin 2024 (https://www.originlab.com/) software and the 3D interaction cluster images were prepared using PyMOL (56) (https://pymol.org/) software.

## Data Availability

The data that support this study are available from the corresponding authors upon reasonable request. The data related to the X-ray structures determined for the apo EstS1 and its complexes have been deposited to the RCSB protein databank under PDB accession codes of PDB ID: (native 1.22 Å), PDB ID: (native 1.5 Å), PDB ID: (Jeffamine), PDB ID: (para-nitrophenol), PDB ID: (para-nitrophenyl butyrate), PDB ID: (Monomethyl phthalate), PDB ID: (Dimethyl phthalate at active site), PDB ID: (Dimethyl phthalate at surface), PDB ID: (diethylhexyl phthalate), PDB ID: (monoethylhexyl phthalate), and PDB ID: (phthalate) respectively.

## Acknowledgment

This work was funded by an industrial grant by THDC INDIA LIMITED. Rishikesh, Uttrakhand, 249201, (Project no. No. THDC/RKSH/ R&D/F-2063/1021) and WTI 2019 grant by the Department of Science and Technology (DST), India (Project No. DST/TMD/EWO/WTI/2K19/EWFH/2019/8 (G)) to P. K. The X-Ray Crystallography facility was provided by Macromolecular Crystallographic Unit, Institute Instrumentation Centre, IIT Roorkee. Ashok Soota Molecular Medicine Facility at Department of Biosciences and Bioengineering, IIT Roorkee was utilized for biochemical experimental work. The computational facility for structure solving was provided by DBT funded project entitled ” Translational and Structural Bioinformatics – BIC at Department of Biotechnology, Indian Institute of Technology, Roorkee (BT/PR40141/BTIS/137/16/2021). SV wants to thank MHRD and KAK wants to thank CSIR for providing fellowship.

