## Supplementary material for "Unveiling Mechanistic and Structural Insights of EstS1 Esterase: A Potent Broad-Spectrum Phthalate Diester Degrading Enzyme"

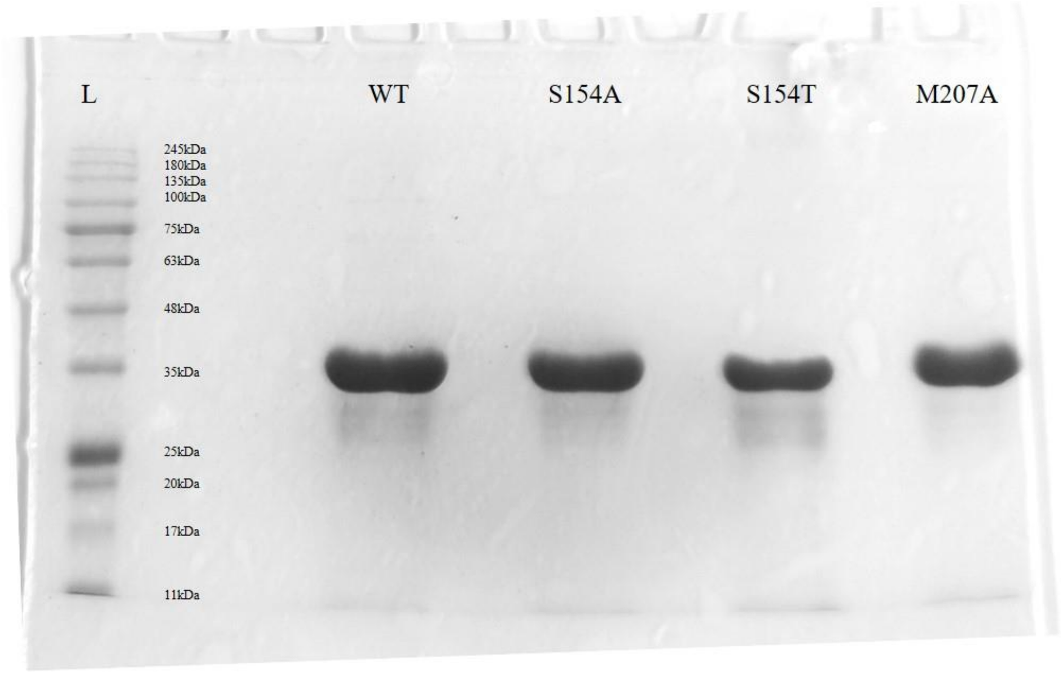


**Figure S1:** An SDS gel image of 6x His-tagged EstS1 wild-type (WT) and S154A, S154T and M207A mutant forms showing clean bands at nearly 34 KDa (L: Protein Ladder)

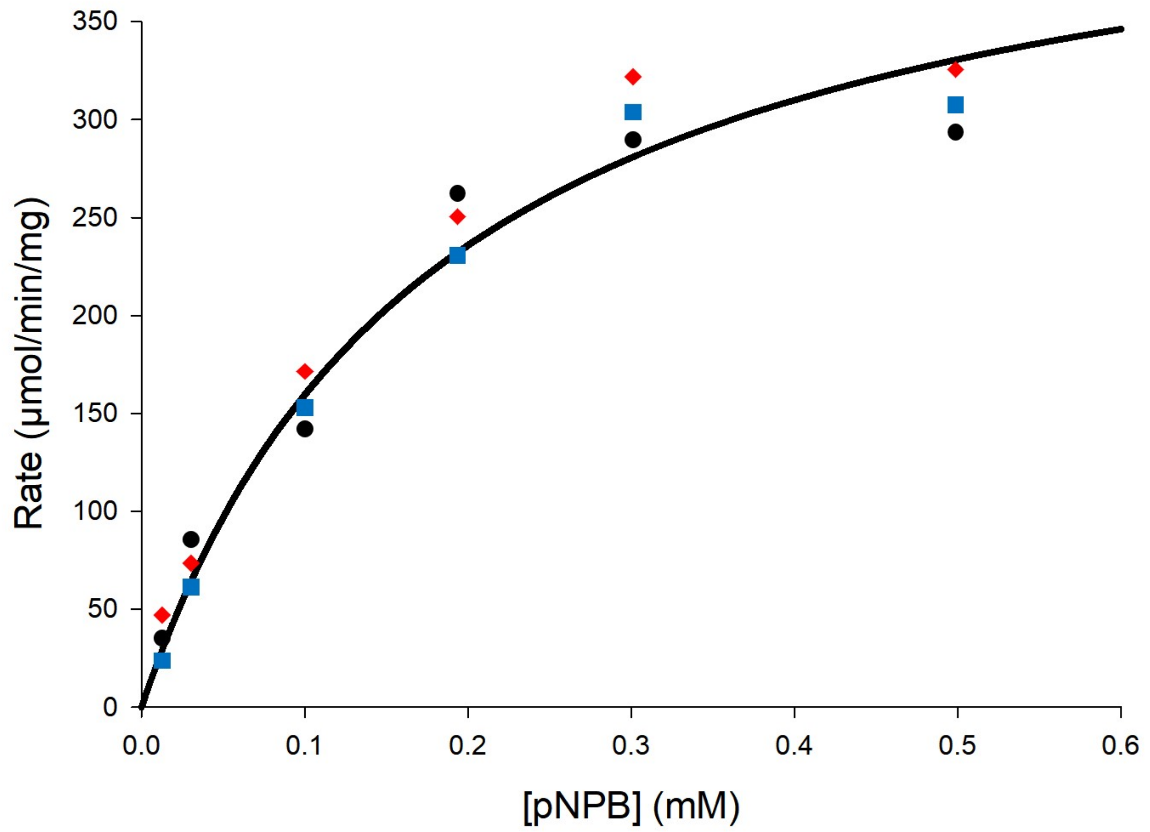


**Figure S2:** Michaelis-Menten curve of EstS1 with para-nitrophenyl butyrate (pNPB) as substrate

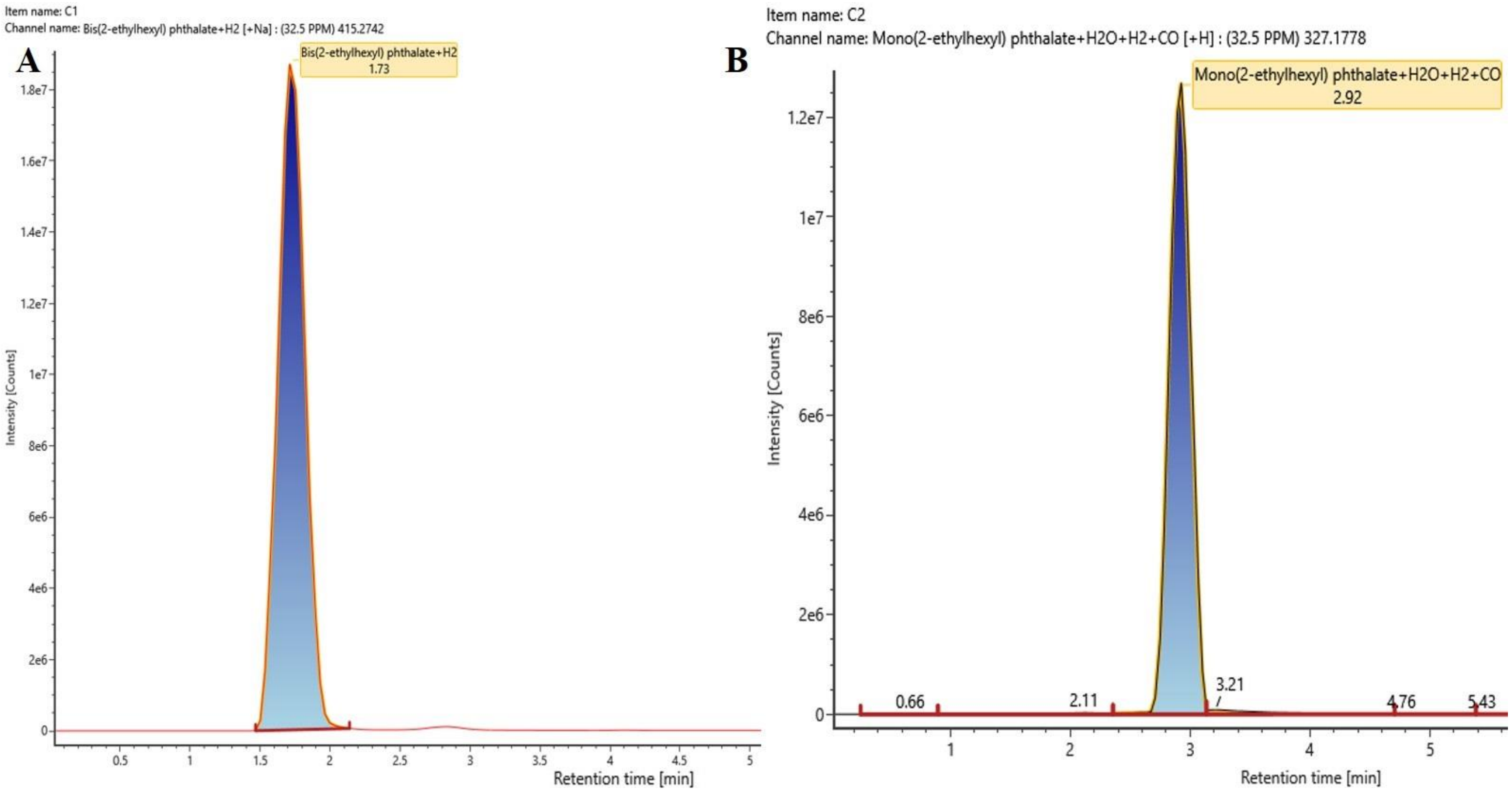


**Figure S3:** Intensity peaks of LC-MS analysis of control samples of **A**) Bis(2-ethylhexyl) phthalate and **B**) Mono(2-ethylhexyl) phthalate displaying the exact retention time of the pure compounds

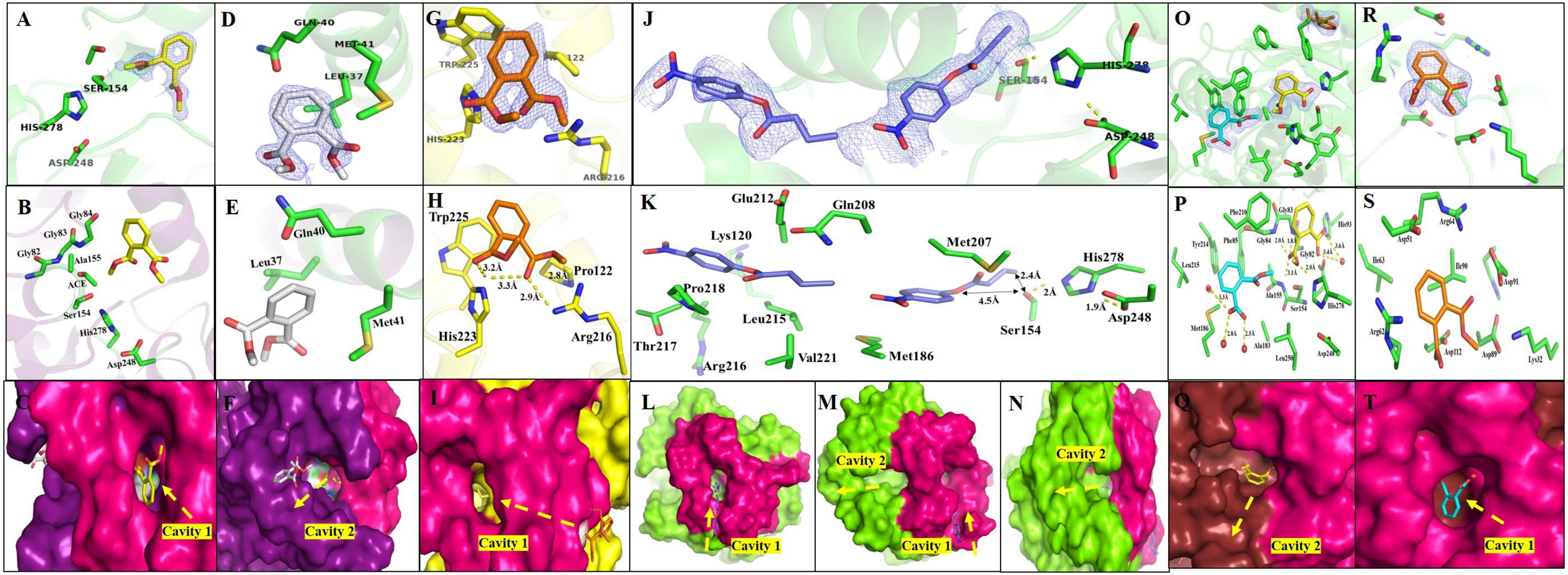


**Figure S4:** High-resolution crystal structures of EstS1 with different substrates and products. A-F) Crystal structure of EstS1-dimethyl phthalate complex at 1.6 Å containing two molecules of dimethyl phthalate. A) Electron density $F_o$-$F_c$ map contoured at 3σ of the first molecule of dimethyl phthalate near the active site. B) Stick representation showing interacting residues of the first molecule of dimethyl phthalate at the active site where acetyl group was also observed at the catalytic Ser154 and hindered the catalysis of dimethyl phthalate even in wildtype EstS1. C) Surface representation of EstS1 showing first molecules of dimethyl phthalate near the active site of EstS1 while entering from cavity 1. D) Electron density $F_o$-$F_c$ map contoured at 3σ of the second molecule of dimethyl phthalate near cavity 2. E) Stick representation showing interacting residues of the second molecule. F) Surface representation of EstS1 showing the second molecule of dimethyl phthalate in complex with EstS1 esterase showing its orientation near cavity 2. G-I) Crystal structure of EstS1 in complex with dimethyl phthalate at 1.5 Å at the outer surface of EstS1 revealing the path followed to enter the catalytic tunnel. G) Electron density $F_o$-$F_c$ map contoured at 3σ of dimethyl phthalate, H) Stick representation showing binding site residues and effective interaction of dimethyl phthalate at the outer surface of EstS1 (yellow dash line marks the bond and bond distances). I) Surface representation of dimethyl phthalate-EstS1 complex showing the relative position of dimethyl phthalate at EstS1 surface showing path followed to reach the entry cavity 1 leading towards the active site. J-N) Crystal structure of EstS1 with para-nitrophenyl butyrate at 1.8 Å. J) Electron density $F_o$-$F_c$ maps contoured at 3σ of para-nitrophenyl butyrate observed at two positions in the EstS1 para-nitrophenyl butyrate complex. K) Stick representation para-nitrophenyl butyrate observed at two positions in the EstS1 showing interaction with active residues and the residues of the entry site. (yellow dash line marks the bond and bond distances) L) Surface representation of para-nitrophenyl butyrate-EstS1 complex displaying first molecule of para-nitrophenyl butyrate at the entry into the catalytic tunnel via cavity 1. M) Surface representation of EstS1 showing first molecule of para-nitrophenyl butyrate at entry site with respect to cavity 1, cavity 2 and the catalytic tunnel. N) Surface representation of EstS1 showing second molecule of para-nitrophenyl butyrate with respect to cavity 2. O-T) Crystal structure of monomethyl phthalate in complex with EstS1 at 2 Å showing electron density of containing 3 molecules of monomethyl phthalate. O) Electron density $F_o$-$F_c$ maps contoured at 3σ of monomethyl phthalate at two positions (cyan colour molecule at active site and yellow colour molecule at the exit site near cavity 2) out of 3 observed in the crystal structure. These two positions accommodated the catalytic tunnel of EstS1 esterase. P) Stick representation of two positions of monomethyl phthalate observed in the crystal structure showing interactions with the active site (cyan colour molecule) and binding site (yellow colour molecule) residues. Q) Surface view of EstS1 showing the second molecule (yellow colour) tending towards cavity 2 or exit site of the catalytic tunnel. R) Electron density $F_o$-$F_c$ map contoured at 3σ of the third molecule (orange colour) of monomethyl phthalate lying at the outer surface of the enzyme. S) Stick representation of the interacting site residues of the third molecule (orange colour) of monomethyl phthalate. T) Surface representation of EstS1 displaying monomethyl phthalate interacting at active site with respect to cavity 1 or entry site. (Yellow dashed arrows represent the direction of entry and exit of the molecule, red sphere represent water molecules and pink colour surface denotes the cap domain).

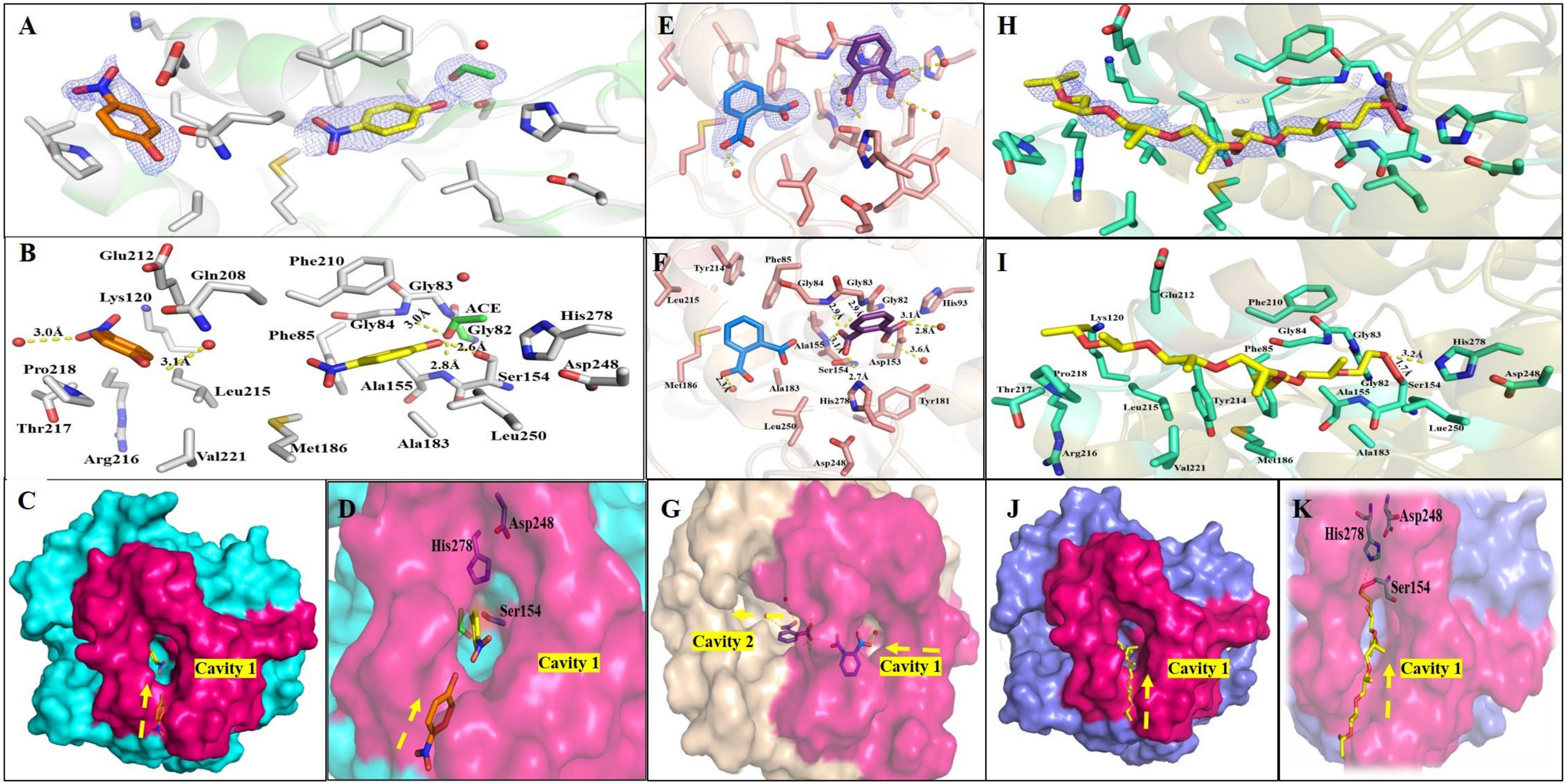


**Figure S5**: Crystal structures of EstS1 with different substrates and products at high resolution. A-D) Crystal structure of para-nitrophenol in complex with EstS1 at 1.5 Å. A) Electron density $F_o$-$F_c$ maps contoured at 3σ of two para-nitrophenol molecules observed in the crystal structure. B) Stick representation of two molecules of para-nitrophenol interacting at the active site (yellow) and a site near cavity 1 (orange) or the entry site. The interacting residues are represented in grey color. C) Surface view of EstS1 displaying the two molecules of para-nitrophenol entering via cavity 1 into the active site of the EstS1 esterase. D) Close-up of surface representation of EstS1 complex with para-nitrophenol displaying the relative position of the two molecules with respect to the active site. E-G) Crystal structure of EstS1 in complex with phthalate at a resolution of 1.5 Å. E) Electron density $F_o$-$F_c$ maps contoured at 3σ of two phthalate molecules observed in the crystal complex structure. F) Stick representation showing interactions of two phthalate molecules positioned at the active site (purple) and in the binding site (blue) with other residues G) Surface view of EstS1 complex with phthalate showing positions of two phthalate molecules in the catalytic tunnel with respect to cavity 1 (entry) and cavity 2 (exit). H-K) Crystal structure of jeffamine in complex with EstS1 at 1.5 Å. H) Electron density $F_o$-$F_c$ maps contoured at 3σ of jeffamine in complex with EstS1 esterase. I) Stick representation displaying the interactions of jeffamine bound at the active site Ser154 of EstS1 esterase, tracing the catalytic tunnel. J) Surface view of EstS1 jeffamine complex showing jeffamine tracing the entry cavity 1 to the active site of the enzyme. K) Surface view of EstS1 jeffamine complex showing jeffamine tracing the catalytic tunnel and interacting at the active site of the enzyme showing its relative position to the active site residues. (Yellow dashed arrows represent the direction of entry and exit of the molecule, red sphere represents water molecules and the pink colour surface denotes the cap domain).

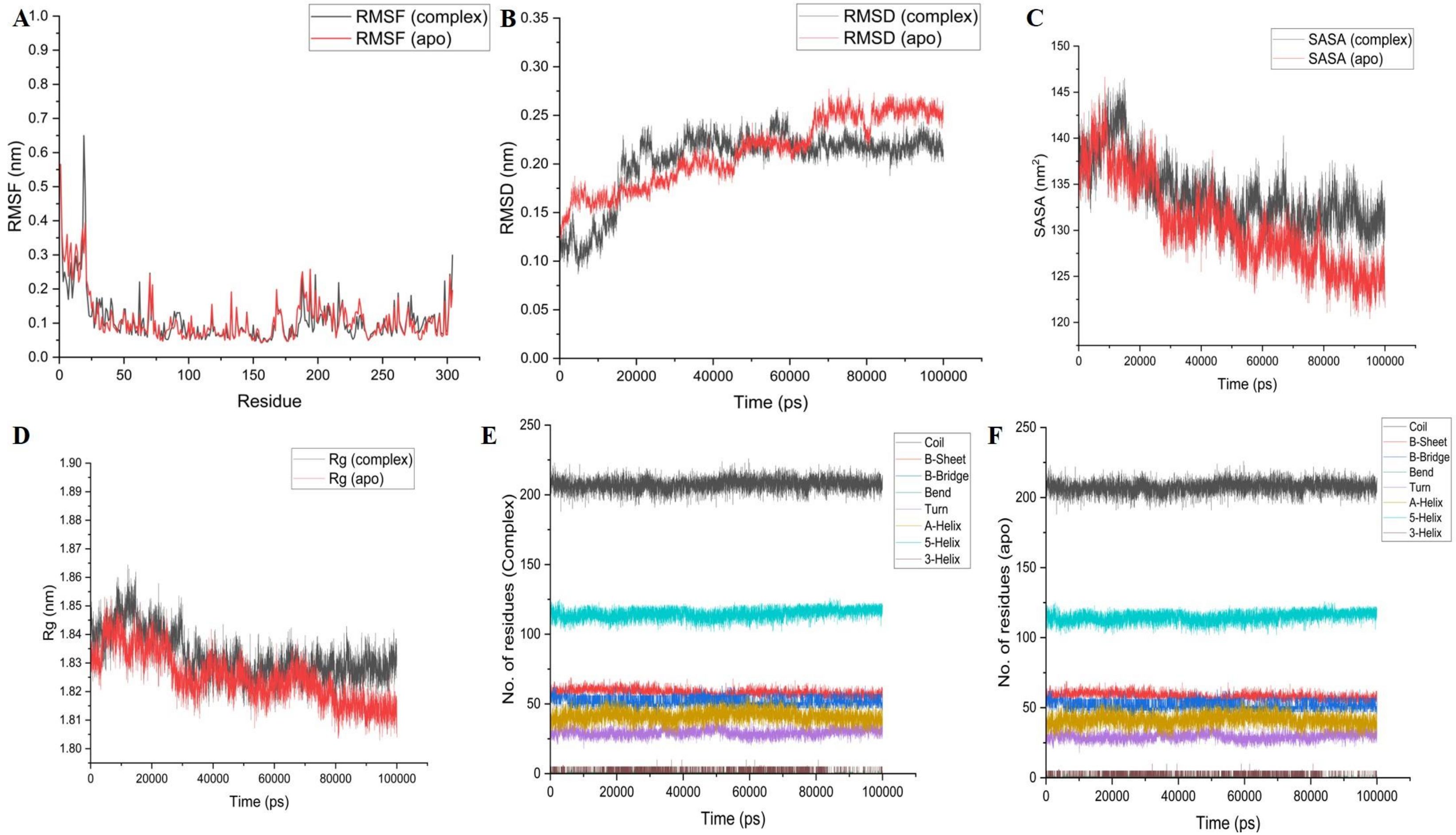

**Figure S6**: Molecular dynamic simulation analysis of the DEHP-EstS1 S154A mutant crystal complex after replacing Ala154 with active site serine residue and the Apo-EstS1. A) Root Mean Square Fluctuation (RMSF) versus time plot of complex and apo-protein. B) Root Mean Square Deviation (RMSD) versus time plot of complex and apo-protein. C) Solvent Accessible Surface Area (SASA) versus times plot of complex and apo-protein. D) Radius of Gyration (Rg) versus time plot of complex and apo-protein. E) Number of residues of EstS1 (in presence of DEHP) contributing to different secondary structures during the trajectory time. F) Number of residues of EstS1 (in absence of DEHP) contributing to different secondary structures during the trajectory time, over the 100ns simulation trajectory

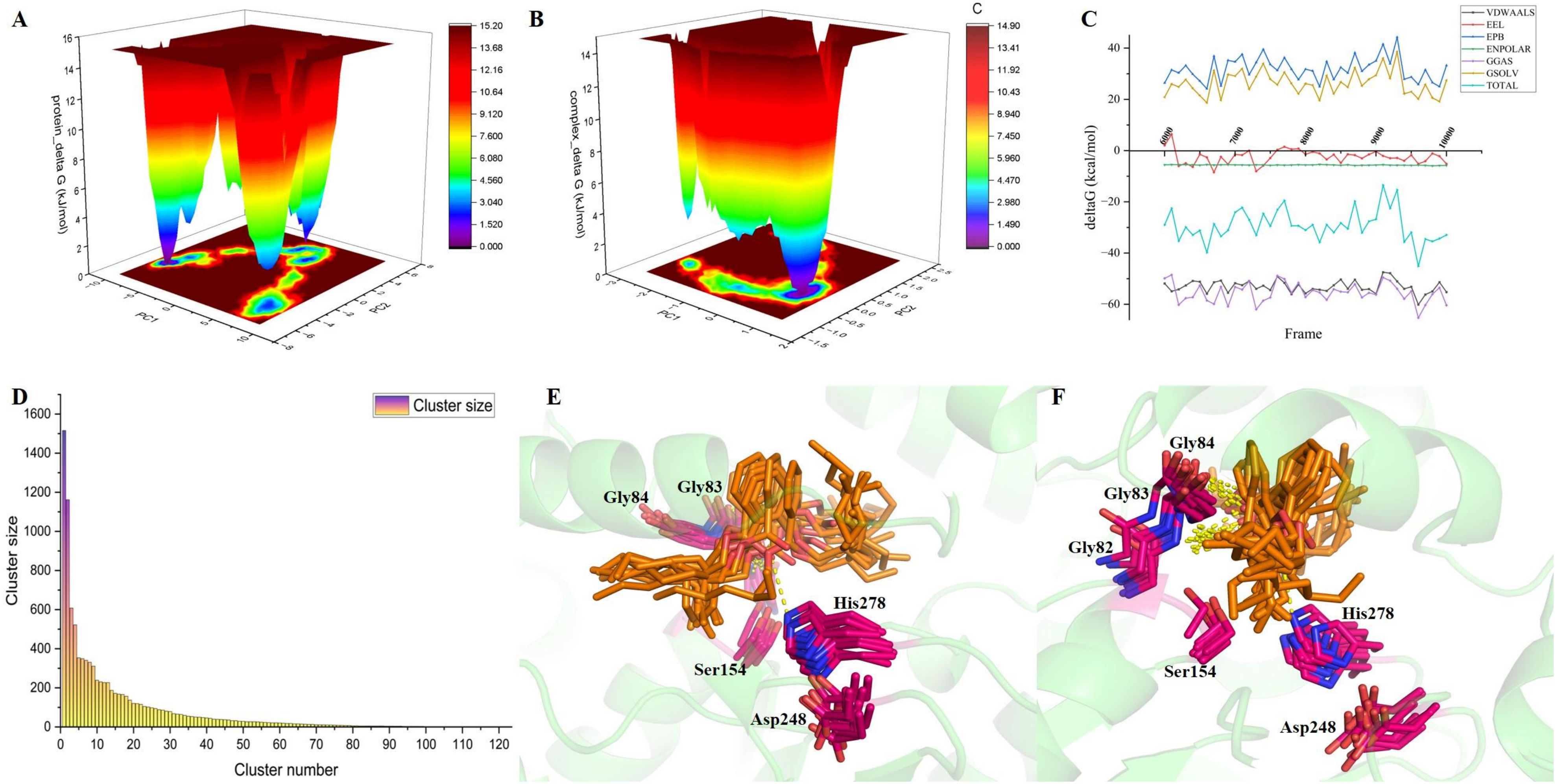

**Figure S7**: Molecular dynamic simulation analysis of the DEHP-EstS1 S154A mutant crystal complex after replacing Ala154 with active site serine residue and the Apo-EstS1. **A)** Free Energy Landscape of apo-EstS1 and **B)** DEHP-EstS1 complex versus principal component 1 and 2 evaluated based on the covariance matrix of different conformational coordinates extracted during the simulation of 100ns. **C)** End state Free binding energy analysis of the DEHP-EstS1 complex for last 40ns of the trajectory using Molecular Mechanics Poisson-Boltzmann Surface Area (MMPBSA) approach. Different components of energy include van der Waals (VDWAALS), electrostatic component of internal energy (EEL), electrostatic contribution to solvation free energy estimated by PB (EPB), non-polar energy (ENPOLAR), DELTA GGAS = Bond + Angle + Dihedral + Electrostatic + van der Waals, DELTA GSOLV = DELTA EPB + DELTA ESURF (proportional to Solvent Accessible Surface Area), TOTAL = DELTA GGAS + DELTA GSOLV. Clustering analysis of all the DEHP-EstS1 complex conformations captured during the simulation trajectory gave **D)** cluster size versus cluster number plot showing that top 10 clusters contain the maximum number of conformations. **E)** Representative conformations of the top 10 clusters displaying interactions with the catalytic triad (Ser154, Asp248 and His278) **F)** Representative conformations of the top 10 clusters displaying interactions with residues (Gly83, Gly84) forming oxyanion hole.

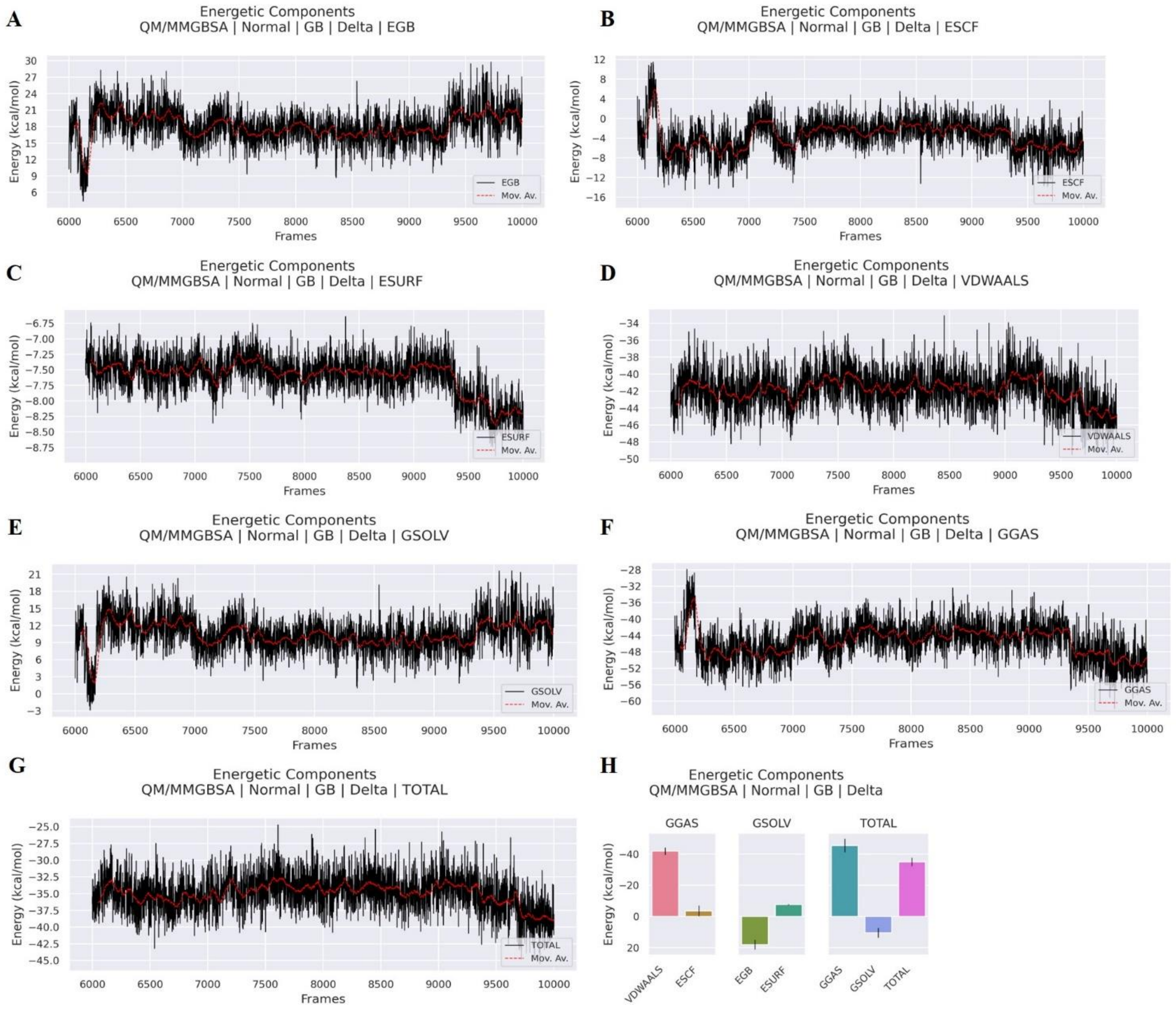


**Figure S8**: QM/MMGBSA analysis for change in different energy components of the DEHP-EstS1 complex interactions over the last 40ns of the simulation trajectory considering Gly82, Gly83, Gly84, Ser154, Ala155, Asp248, His278 and DEHP residues in the quantum mechanical region. A) Electrostatic contribution to the solvation-free energy calculated by Generalized Born model (EGB) B) Energy of self-consistent field (ESCF) considering QM and QM/MM region C) Non-polar component of the solvation energy (ESURF) D) Energy contribution by van der Waals

(VDWAALS) interaction E) Delta GSOLV = Delta EGB + Delta ESURF F) Delta GGAS = Delta VDWAALS + Delta ESCF G) TOTAL = Delta GGAS + Delta GSOLV H) Bar plot showing average change in all the energy components contributing for interactions in the QM and QM/MM system during the simulation trajectory

**Table S1**: Primers used for site-directed mutagenesis

| **Mutant** | **Primers** |
|---|---|
| S154A | FP: GTGGCCGGGGACGCGGCCGGCGGCAATTTG |
| | RP: GCCGCCGGCCGCGTCCCCGGCCACCACAATC |
| S154T | FP: GATTGTGGTGGCCGGGGACACCGCCGGCGGCAAT |
| | RP: CGCCAAATTGCCGCCGGCGGTGTCCCCGGCCAC |
| M207A | FP: TTTTGGAGGCGGACGCGGCGCAATGGTTTGGCGAACAATA |
| | RP: GCCAAACCATTGCGCCGCGTCCGCCTCCAAAAGATAA |
